# Depth-dependent scaling of axial distances in light microscopy

**DOI:** 10.1101/2024.01.31.578242

**Authors:** S.V. Loginov, D.B. Boltje, M.N.F. Hensgens, J.P. Hoogenboom, E.B. van der Wee

**Affiliations:** Department of Imaging Physics, Delft University of Technology, Delft, The Netherlands; Delmic B.V., Delft, The Netherlands

## Abstract

In volume fluorescence microscopy, refractive index matching is essential to minimize aberrations. There are however, common imaging scenarios, where a refractive index mismatch (RIM) between immersion and sample medium cannot be avoided. This RIM leads to an axial deformation in the acquired image data. Over the years, different axial scaling factors have been proposed to correct for this deformation. While some reports have suggested a *depth-dependent* axial deformation, so far none of the scaling theories has accounted for a depth-dependent, non-linear scaling. Here, we derive an analytical theory based on determining the leading constructive interference band in the objective lens pupil under RIM. We then use this to calculate a depth-dependent re-scaling factor as a function of the numerical aperture (NA), the refractive indices *n*_1_ and *n*_2_, and the wavelength *λ*. We compare our theoretical results with wave-optics calculations and experimental results obtained using a novel measurement scheme for different values of NA and RIM. As a benchmark, we recorded multiple datasets in different RIM conditions, and corrected these using our depth-dependent axial scaling theory. Finally, we present an online web applet that visualizes the depth-dependent axial re-scaling for specific optical setups. In addition, we provide software which will help microscopists to correctly re-scale the axial dimension in their imaging data when working under RIM.

## 1. Introduction

Optical sectioning has enabled imaging of large volumes by fluorescence microscope (FM), as realized in confocal, two-photon microscopy [1], structured illumination microscopy (SIM) [2] and light sheet microscopy [3]. For optimal imaging, aberrations need to be minimized by avoiding a RIM between the detection microscope objective immersion medium *n*_1_ and specimen *n*_2_ [1]. Failing to do so results in the blurring of the point spread function (PSF) of the microscope and therefore a loss in resolving power, as well as in a deformation of the recorded volume along the optical axis.

This axial deformation arises from the refraction of the peripheral rays on the RIM interface, causing an axial shift of their focal point with respect to the focal point of the paraxial rays [4]. This effect can be characterized using focal shift Δ *f* = AFP − NFP, where AFP is the actual focal position (the real position of the object) and NFP is the apparent or nominal focal position (the microscope z-position where the object is found in focus) [1]. Just as the true lateral distances are recalculated back from an image using the magnification of the objective *M*, so ought the AFPs (i.e. the true axial distances) to be re-scaled using the re-scaling factor ζ = AFP / NFP. The accurate knowledge of the re-scaling factor enables reliable quantitative volumetric microscopy.

While it is generally unfavorable to have a refractive index mismatch, for e.g. illumination intensities in confocal microscopy, there are common imaging scenarios in which a mismatch is still present in the most optimal configuration [1, 4]. For instance, as the resolving power of the microscope depends on the NA of the detection objective, high-NA oil immersion objectives (*n*_1_ = 1.52) are used to image water-like specimens (*n*_2_≈ 1.33). In addition, embedding and fixation media rarely match exactly, in terms of refractive index, the immersion media of air, water, silicone, glycerol or oil objectives, leading to the axial deformation of the imaged volumes.

Refractive index mismatches are often more pronounced in integrated cryogenic fluorescence and/or correlative microscopy [5–8]. In such systems, light and electron microscopy are combined in a single setup, and air objectives are often used for imaging specimens with a higher refractive index. The use of air objectives is a straight-forward choice, as the specimen resides in a vacuum chamber for electron microscopy. In cryogenic fluorescence microscopy the specimen is cooled to temperatures below 120 K, which makes the use of a non-touching air objective favorable, as any other (touching) immersion objectives are challenging in terms of engineering [9–12].

The development of the confocal microscope, along with its optical sectioning capability, allowed for 3D imaging, and hence the need for axial scaling theories arose. In the paraxial approximation, the axial distances are simply re-scaled using the ratio of the two refractive indices [13], which works well for low NA objectives. Visser *et al*. presented a scaling theory which is based on the contribution of the high-angle (or marginal) rays to the axial scaling [14]. More recently, several scaling theories have been presented, which result in re-scaling factors somewhere in between the paraxial and high-angle approximations [4, 15, 16]. The most accurate method to determine axial scaling are full wave-optics calculations of the microscope’s point spread function under RIM [17–19]. As these calculations are computationally expensive and complex, they are hardly used by microscopists to calculate the axial re-scaling factor.

The re-scaling factor can also be measured experimentally by observing the NFP of a fluorescent bead or interface through a medium with refractive index *n*_2_, whilst knowing the AFP of said bead or interface. This can be done by constructing a sample cell where the fluorescence is present far from the coverslip, which is later filled with a liquid with *n*_2_ [4, 20, 21]. The AFP can be obtained through imaging with an objective where *n*_1_ = *n*_2_ [4, 20, 21] or by measuring Fabry–Pérot fringes in the transmission spectrum of the cell [21]. Alternatively, the apparent axial deformation of a spherical object, larger than the PSF of the microscope, can be used to measure the axial scaling [14, 22]. Recently, a different approach was presented where a coverslip was step-wise coated with a low-index polymer (*n*_2_ ≈ 1.33) and the AFP was measured using stylus profilometry [23]. With this method, the axial scaling can be measured in the range of a few microns from the coverslip, while the former methods are used to measure tens of microns away from the coverslip.

While the explicit axial scaling theories in literature are all depth-independent, there exist some reports in literature that this factor is actually depth-dependent. In 1993, Hell *et al*. wrote *“*…*it can be expected that the regions close to the cover glass are slightly more scaled than those in deeper regions of the specimen*.*”* [17]. Later, wave-optics calculations showed a non-linear dependence of the focal shift Δ *f* on the imaging depth for high-NA objectives and large RIMs [18, 19]. For instance, Sheppard and Török reported a non-linear dependence of the focal shift Δ *f* at a depth <30 µm from the coverslip using wave-optics calculations (NA = 1.3, *n*_1_ = 1.52, *n*_2_ = 1.33) [18], where the re-scaling factor was 5% larger close to the coverslip than at large distances. More recently, the measurements by Petrov *et al*. showed significant non-linear axial scaling for imaging depths < 4 µm [23].

There is, up to now, no straightforward equation which can be used to explicitly calculate the re-scaling factor as a function of depth. Moreover, a depth-dependence of the re-scaling factor has not been measured experimentally for large depths, and for several NAs and multiple RIM conditions.

Here, we present an analytical theory which calculates the depth-dependent re-scaling factor as a function of the NA, the refractive indices *n*_1_ and *n*_2_, and the wavelength *λ*. The gist of the theory is in the determination of the leading constructive interference band in the objective lens’ pupil under RIM. We compare the theory to both full wave-optics calculations and experiments.

In the experiments, we have imaged the gap between two substrates that were brought closer to each other step by step. By filling the space between the substrates with a liquid with index *n*_2_ we were able to measure the NFP, while the AFP was determined independently from the microscope by the piezo-stage holding the top substrate. We have measured the re-scaling factor ζ for several objectives with various NAs, immersion refractive index *n*_1_, and sample refractive index *n*_2_, with both RIMs where *n*_1_ < *n*_2_ and *n*_1_ > *n*_2_ in a wide range of depths and compared them to the analytical theory and the wave-optics calculations. We demonstrate that the axial re-scaling of 3D microscopy data, recorded with a refractive index mismatch, using the depth-dependent re-scaling factor outperforms the re-scaling using existing linear re-scaling theories from literature. Finally, we provide the reader with an online web applet where one can visualize the depth-dependent axial re-scaling factor for their specific optical setup and Python software to re-scale data acquired under RIM.

## 2. Scaling of axial distances due to refractive index mismatch

### 2.1. Geometrical optics

Given a fluorescent object emitting light in an ideal, spherical fashion, a flat interface with a refractive index mismatch will cause disturbance to the fluorescent light propagation – in the form of refraction. When an objective lens is collecting light under RIM, this effect will be more pronounced with increasing NA of the objective lens (OL) (and therefore collection angle), see Figure 1a for the overall geometry.

**Fig. 1.**
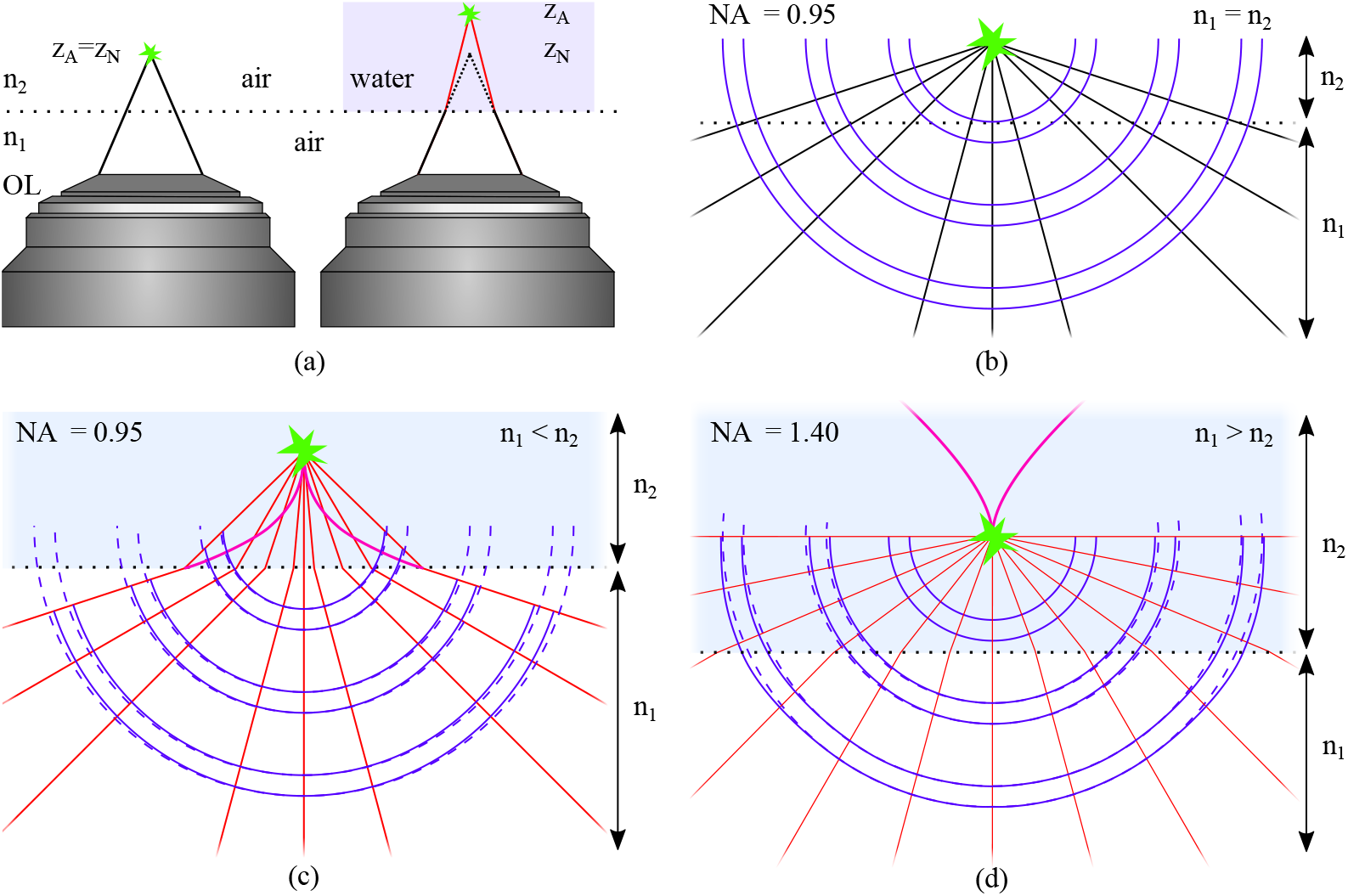
The effects of RIM demonstrated with geometrical optics. **(a)** An OL imaging a fluorescent object (green star) with- and without refractive index mismatch, left and right respectively. The dotted boxes are detailed out in panels b through d. **(b)** A fluorescent object (green star) sits *z* _*A*_ = 10 µm deep from the refractive index interface (dotted line) between *n*_1_ and *n*_2_. With *n*_1_ = *n*_2_ = 1.0, rays (black) and wave-fronts (blue) are depicted. **(c)** In the case where *n*_1_ = 1.0, *n*_2_ = 1.33 the ideal spherical wave-fronts (dashed blue) are deformed (solid blue) after crossing the refractive index (RI) interface. Rays are depicted in red to indicate RIM and the caustical surface (purple) shows where geometrical optics breaks down. **(d)** In the opposite RIM case, where *n*_1_ = 1.52, *n*_2_ = 1.33, again the wave-fronts are deformed under RIM and total internal reflection occurs. The NA is 0.95 **(b, c)** and 1.4 **(d)**.

This is further illustrated using geometrical optics in Figure 1b through d, where a light source is located 10 µm away from the interface between *n*_1_ and *n*_2_. In Figure 1b no RIM is present (*n*_1_ = *n*_2_ = 1.0), where the undisturbed wave-fronts and rays are shown in solid blue and black respectively. The dotted horizontal line depicts the interface with the RIM. Figure 1c and d show two cases where RIMs are present respectively for *n*_1_ < *n*_2_ and *n*_1_ > *n*_2_. The black rays follow Snell’s law when crossing the interface, resulting in the wave-fronts (solid blue) deviating from the ideal spherical shape (dashed blue). The caustic surface (purple) indicates where geometrical optics breaks down. From this simple illustration, we see that the effect of RIM becomes more pronounced with increasing collection angles (numerical aperture) and increasing RIM contrast Δ*n* = |*n*_2_ − *n*_1_|.

When imaging with a microscope, there are two (independent) axial coordinates to be considered: the actual depth of the fluorescent emitter *z* _*A*_, as measured from the RIM interface (*z* _*A*_ = 10 µm in Figure 1) and the nominal depth of the focal plane of the microscope into the sample *z*_*N*_. A 3D image stack recorded with a microscope under RIM has *z*_*N*_ as an axial coordinate. Similar to using the magnification *M* to re-calculate the true lateral dimensions, the axial coordinates should be re-scaled using the re-scaling factor ζ≡ *z* _*A*_ / *z*_*N*_ to obtain a 3D stack with the actual depth *z* _*A*_ as an axial coordinate.

Geometrical optics can produce several estimates for the re-scaling factor ζ. The paraxial rays are nicely focused, even under RIM, producing the estimate [13]:

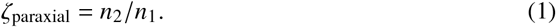

Another geometrical optics estimate comes from Visser *et al*., obtained from the marginal rays still fitting into the NA [14]:

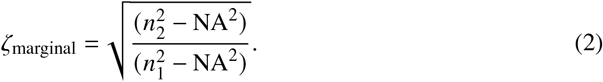

The two re-scaling factor values ζ_paraxial_ and ζ_marginal_ are irreconcilable in a modern high-performance objective lenses (with NA → 0.95 × *n*_1_). For the example shown in Figure 1c, ζ_paraxial_ = 1.33 and ζ_marginal_ = 2.98, whilst for Figure 1d ζ_paraxial_ = 0.88 and ζ_marginal_ → 0.

Although these estimates are derived from geometrical optics, ζ_paraxial_ and ζ_marginal_ do provide lower and upper bounds when *n*_1_ < *n*_2_ (reverse when *n*_1_ > *n*_2_) and are still useful checks in the wave-optics treatment.

### 2.2. Analytical expression describing depth-dependent axial scaling

When considering the PSF of a wide-field microscope under RIM two intrinsic length scales need to be considered: (i) several strictly geometrical parameters *n*_1_, *n*_2_, *z* _*A*_, *z*_*N*_, & NA, and (ii) the physical parameter of the (vacuum) wavelength of light used *λ* = 2π / *k* which is independent from the exact RIM geometry, where *k* is the wave number. With changing imaging depth, the geometrical parameters *z* _*A*_, *z*_*N*_ do change, while the wavelength of light *λ* does not. Following the derivation outlined below, we find it is exactly the interplay between the two length scales that yields a depth-dependent re-scaling factor:

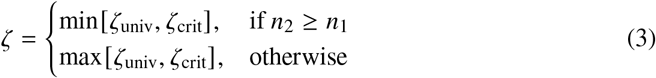

with

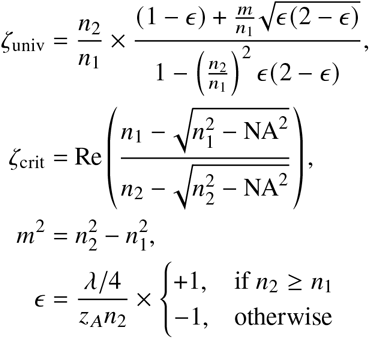

where ζ is the axial re-scaling factor (ζ *≡z* _*A*_ / *z*_*N*_), *n*_1_ is the immersion refractive index, *n*_2_ the sample refractive index, *λ* the wavelength (in vacuum), *z* _*A*_ the actual depth of the imaged object and NA the numerical aperture. The re-scaling factor ζ is a combination of the depth-dependent and NA-independent ζ_univ_ which approaches the paraxial limit ζ_paraxial_ for large depths, and the depth-independent and NA-dependent ζ_crit_ at shallow depths. ζ_crit_ becomes relevant at such shallow depths where the assumptions used to derive ζ_univ_ breakdown (see for details section 2.4).

### 2.3. Derivation of analytical theory through wave-optics

With geometrical optics ambiguous in defining the re-scaling factor ζ we have to turn to wave-optics. The most complete wave-optics theory describing aberrations occurring under RIM was developed by Hell *et al*. [17]. Their calculation scheme takes into account the vectorial nature of light and, is in essence, equivalent to the 3-fold application of the scalar model described in [24]. The scalar model in turn is based on the general Diffraction Integral applied to the ray fans in RIM [25–27]. A scalar (radially symmetric) PSF of a wide-field microscope under RIM is an integral over Bessel-beams:

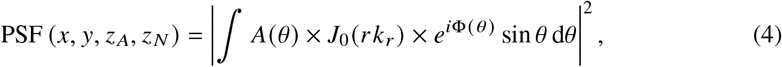

where the PSF is dependent on both the actual depth *z*_*A*_ and on the nominal depth *z*_*N*_, as an image with intensities *I* (*x, y, z*_*N*_) is a convolution of fluorophore distribution *f* (*x, y, z* _*A*_) and the PSF(*x, y, z* _*A*_, *z*_*N*_). Furthermore, 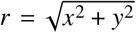 as the PSF is cylindrically symmetric, the amplitude factor 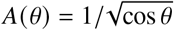 is due to the Abbe-sine condition, *J*_0_ is the Bessel function, *i*^2^ = −1 and the integral is taken over the NA of the objective lens. The effects of RIM are engraved in the phase function Φ(*θ*):

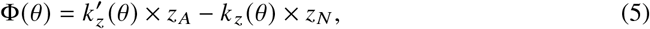

where *k*_*z*_ and 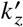 are the axial wave-numbers of the Bessel beams, respectively for a matched case and one undergoing RIM. In the full vectorial treatment, there are three integrals of the form of Equation 4; *I*_0_, *I*_1_ and *I*_2_ (one for each of the E-vector directions, with different orders of Bessel functions). The first integral *I*_0_ has a Bessel function of the order zero and is, in essence, the Diffraction Integral used in the scalar-light theory. The other two integrals *I*_1_ and *I*_2_ of the full vectorial theory contain the same phase function Φ *θ* as in *I*_0_, and, moreover, they vanish at *r* = 0. Thus, we will consider only the scalar-light version of the wave-optics Equation 4 for finding an explicit analytical form of the re-scaling factor ζ in the following paragraphs.

Compared to the ambiguous behavior of geometrical optics, Equation 4 provides a clear prediction of the behavior of a microscope imaging under RIM. For a given depth *z* _*A*_ of a fluorescent object there is a focal plane depth *z*_*N*_ (*z*_*A*_) where the PSF is maximal (i.e. the object is in focus), and ζ (*z* _*A*_) *≡z* _*A*_ / *z*_*N*_ (*z* _*A*_). Under RIM, two intrinsic length scales need to be considered: (i) several strictly geometrical parameters *n*_1_, *n*_2_, *z* _*A*_, *z*_*N*_, and NA. (ii) the (vacuum) wavelength *λ* = 2π / *k* which is totally independent from the RIM geometry. These two length scales are fused together in Equation 4 via the phase function Φ (*θ*). With changing imaging depth, the geometrical parameters also change (e.g. the path length difference between the paraxial and the marginal rays). Reversely, the wavelength of light does not depend on the depth. So it is the interplay between geometrical and physical length scales that results in a depth-dependent re-scaling factor ζ (*z* _*A*_) *≠* const.

To find ζ without calculating the integrals like Equation 4 we estimate the maximum of such an integral by finding the leading terms of a simpler integral:

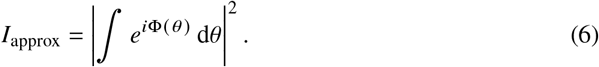

To estimate the maxima positions of *I*_approx_ and assuming that *e*^*i*Φ^ is oscillating fast (for example at large depth), we try to find the stationary points *θ*^*∗*^ where dΦ / d*θ* = 0 [28]. The structure of Φ (*θ*) in case of *n*_1_ < *n*_2_ is shown in Figure 2. The axial wave numbers under RIM and without RIM, 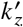 and *k* _*z*_ belong to the Ewalds’ spheres (Figure 2a, red and black), with 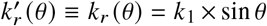 conserved under RIM due to Snell’s law. Radial wave number conservation allows the use of sin *θ* d*θ* = d*k*_*r*_ / *k*_2_, making Equation 4 more similar to *I*_approx_. The two integrals (Equation 4 and Equation 6) are taken over spherical segments having the same width, but different radii due to RIM, where the larger sphere is flattened compared to the smaller one. The RIM phase-function Φ (*θ*) re-scales these spheres in the axial directions in *z* _*A*_ and *z*_*N*_, see Figure 2b. The stationary points *θ*^*∗*^ are found where the two re-scaled spheres are running in parallel to each other (dashed lines, dΦ / d*θ* = 0). There is always a stationary point at *θ* = 0, and a stationary point might appear for a single non-zero *θ*. For a given *z* _*A*_ and *z*_*N*_, there is an interference between the contributions from the two critical points (*e*^*i*Φ(0)^ and *e*^*i*Φ(*θ ∗*^)) which controls the shape of the PSF. And when the interference is the most constructive – then the object appears in focus.

**Fig. 2.**
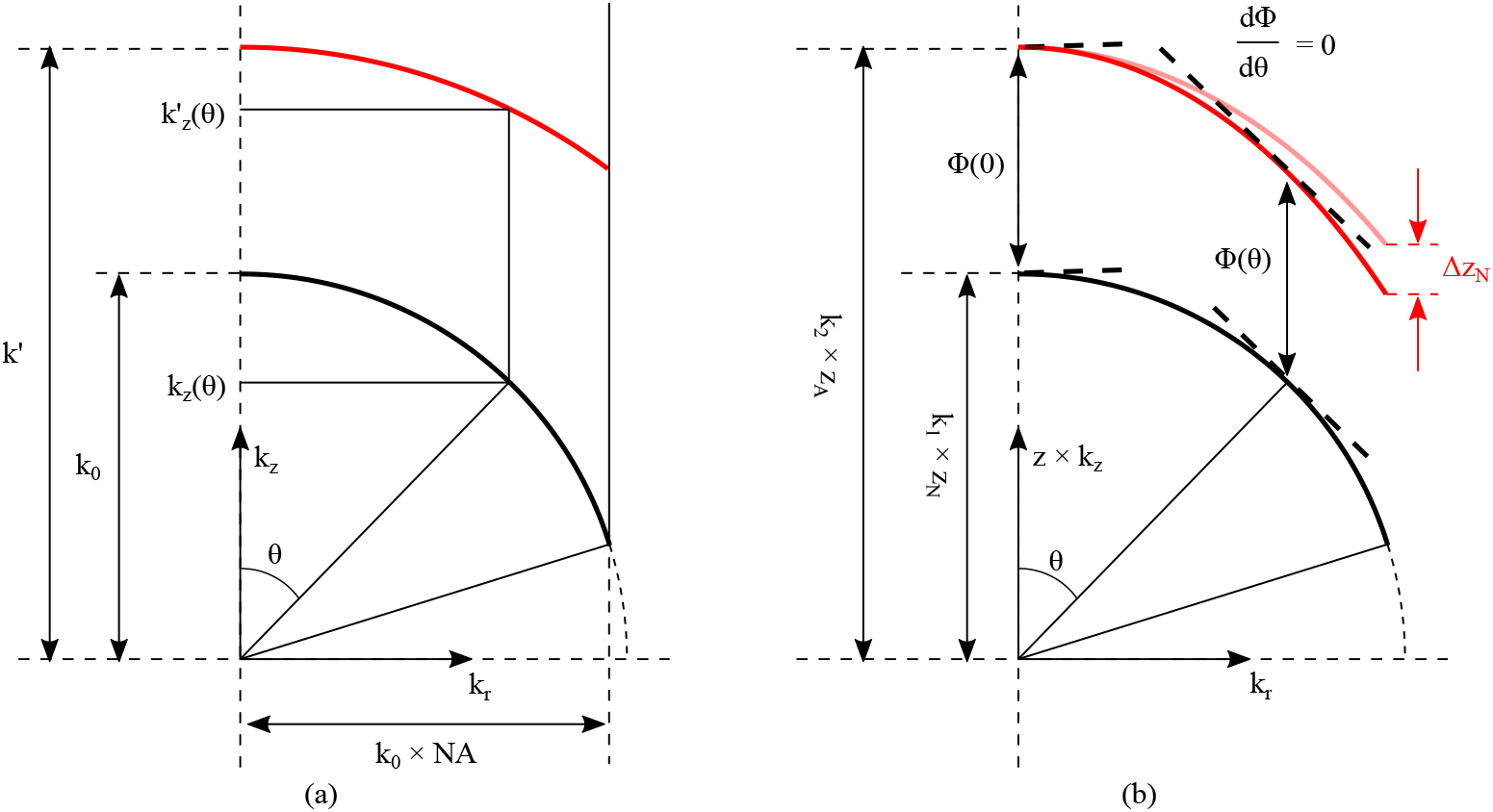
The structure of Φ *θ* for *n*_1_ < *n*_2_, where the diffraction integral in k-space is illustrated with the arcs of Ewalds’ spheres. *k*^′*z*^ and *k* _*z*_ are the axial wave numbers with and without RIM. **(a)** After transition to a new medium the NA is preserved (due to the Snell’s law) while a different Ewalds’ sphere is used, going from black (no RIM) to red (under RIM). **(b)** The RIM phase-function Φ *θ* re-scales (multiplies) the spheres in the axial directions in *z* _*A*_ (red) and *z*_*N*_ (black), and the phase derivative can be set to zero only at one point on the arc depending on the exact values of *z*_*N*_ and *z* _*A*_ (for non-zero *θ*). The Ewald sphere curvature under RIM changes with changing depth (red translucent, Δ*z*_*N*_) and with it the stationary point dΦ/d*θ* = 0.

To find the non-zero stationary point *θ*^*∗*^ we rewrite Φ using Ewalds’ spheres and Snell’s law:

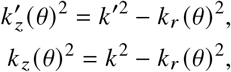

so that when differentiating Φ by *k*_*r*_ :

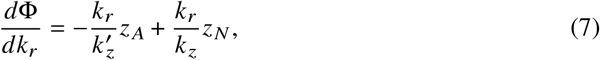

and at a stationary point 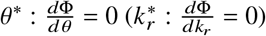

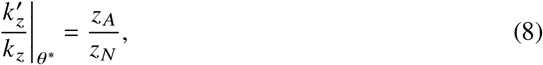

that is the larger radius and thus flatter spherical segment, was multiplied by the larger depth *z*_*A*_ and the smaller radius and more curved spherical segment by the smaller *z*_*N*_ (respectively red, and black in Figure 2). The distance in the vertically stretched segments is the phase function Φ (*θ*) as sketched in Figure 2b. The single intermediate stationary point has to be a minimum for *n*_1_ < *n*_2_, as the difference 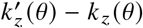 in Figure 2a increases for larger angles *θ* (radial wavenumbers *k*_*r*_) than it is at *θ* = 0 (*k*_*r*_ = 0):

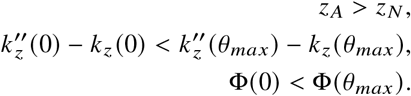

The stationary phase points of the integral in Equation 6 correspond to the rays of geometrical optics [26–28]. Hence, the re-scaling factor must fall within the bounds set by ζ_paraxial_ and ζ_marginal_. Contrary to the purely geometrical optics approach, we have now two contributions to balance: the paraxial one *e*^*i*Φ(0)^ and the intermediate one *e*^*i*Φ(*θ ∗*^).

We estimate for which *z* _*A*_ / *z*_*N*_ the two stationary point contributions *e*^*i*Φ(0)^ and *e*^*i*Φ(*θ ∗*^), build the largest constructive interference. The relative strengths of the two leading contributions in the integral *I*_approx_ (Equation 6) depend on the width of the stationary points of Φ, and hence on the second derivative of Φ. The contributions of stationary-points in Equation 4 are also weighted by pupil-amplitude factor *A* (*θ*). Instead of finding the exact weight of each contribution, we study how the phase function Φ (*θ*) behaves in several focal positions *z*_*N*_ (*z*_*A*_), which can be determined by full vectorial PSF calculations. This is shown in Figure 3a for NA = 0.95, *n*_1_ < *n*_2_. The phase function at the second stationary point lies between 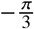 and 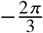 from the phase function at the first stationary point. Therefore, we select the following condition when deriving an explicit formula for ζ :

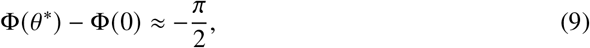

where a fixed −π / 2 separation is assumed, as if the two stationary points constitute a single and uninterrupted constructive interference area when the microscope is in focus. With this approximate condition set, Snell’s law, and the Ewalds’ spheres scaling Equation 8 established, we solve a quadratic equation to derive an explicit formula for ζ. From Equation 9 and the definition of ζ we have:

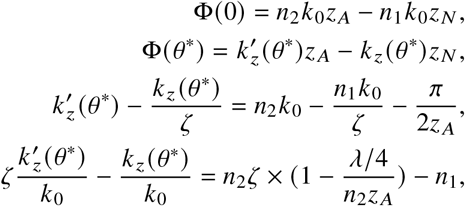

where the dimensionless parameter 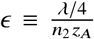 appears. This parameter embodies the before-mentioned interplay between the purely geometrical and purely physical length scales of the RIM. Applying Equation 8:

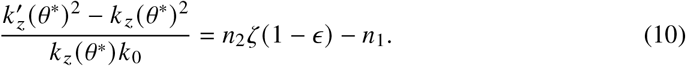

**Fig. 3.**
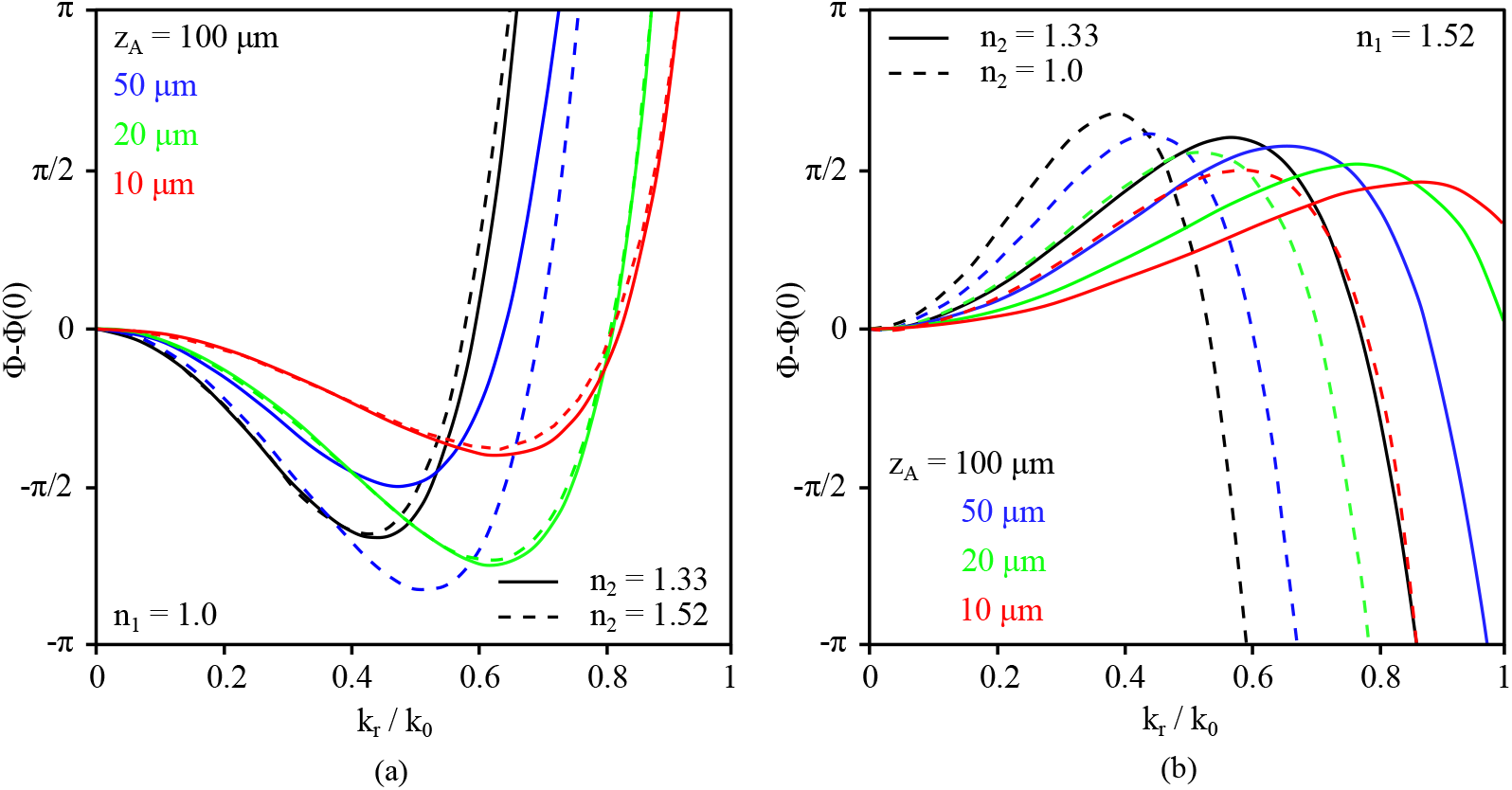
The RIM phase function difference Φ − Φ (0) taken in the focal position *z*_*N*_ (*z*_*A*_), determined from the wave-optics calculations for fluorophore depths *z* _*A*_ set to 10 µm (red), 20 µm (green), 50 µm (blue) and 100 µm (black). **(a)** Objective NA of 0.95, *n*_1_ = 1.0, *n*_2_ = 1.33 (solid lines) and *n*_2_ = 1.52 (dashed lines). **(b)** Objective NA of 1.4, *n*_1_ = 1.52, *n*_2_ = 1.33 (solid lines) and *n*_2_ = 1.0 (dashed lines). For both cases, the second stationary point is somewhere between π/3 and 2π/3 from the paraxial stationary point at *k*_*r*_ /*k*_0_ = 0.

Using Ewalds’ spheres, Snell’s law, and Equation 8 yields:

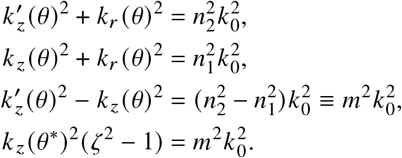

Going back to the left hand-side of Equation 10:

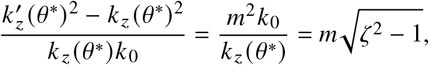

and hence for the equation:

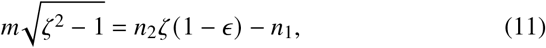

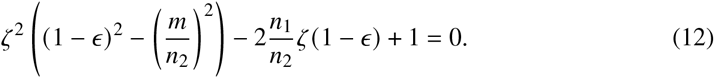

Which, when solved for ζ leads to:

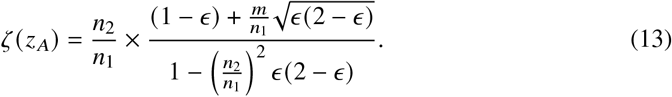

When imaging under the RIM condition *n*_1_ > *n*_2_, the derivation is similar, with some slight differences. The red and black colors in Figure 2 should be switched, which changes the second stationary point from a minimum to a maximum:

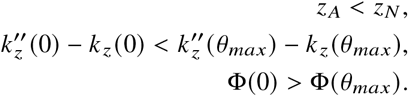

as is shown in Figure 3b where the second stationary point is observed between + π / 3 to + 2π / 3 from the first. With a fixed separation of + π / 2 assumed, we start the derivation with a reversed sign on the right-hand side of Equation 9 and thus obtain a reversed sign of *ϵ*. Together with *m*^2^ < 0 the same formula Equation 13 therefore remains valid for the case of *n*_1_ > *n*_2_.

### 2.4. Critical value at shallow depths

At very large depths, the re-scaling factor ζ approaches the paraxial regime (Equation 1), as *∈* → 0. At very shallow depths (i.e. large *ε*) Equation 13 becomes problematic: ζ (*z* _*A*_ = 0) = ∞ when *n*_2_ > *n*_1_ and ζ (*z*_*A*_ = 0) = 0 for *n*_2_ < *n*_1_, going beyond the bounds of the geometrical optics (ζ_marginal_ in Equation 2). This is related to the fact that the actual value of the NA of the objective lens was omitted in the derivation so far.

At ever decreasing depths, the second critical point *θ*^*∗*^ moves to higher wave numbers *k*_*r*_ and higher angles *θ* (peak of the phase difference in Figure 3) until it falls outside the NA of the optical system (Figure 2a). Hence, there is some (NA dependent) depth below which the assumptions we used to derive Equation 13 break down. We will hence derive a NA dependent critical value of re-scaling ζ_crit_ to be used at small depths.

With the whole range of phase function values Φ (*θ*) becoming smaller than π / 2 at small depths, the earlier condition on the separation of the critical points (Equation 9) can not be satisfied anymore within the confines of the NA. At such small depth, the entire pupil participates in constructive interference and the breakdown condition can be written as:

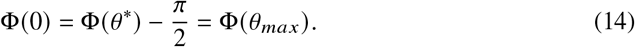

From the definition of Φ(*θ*) and *k*_*r*_ (*θ*_*max*_) = *k*_0_ × NA:

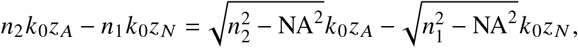

And as 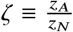 one gets:

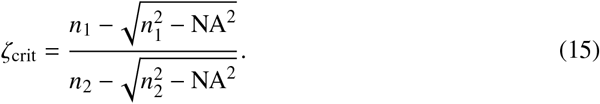

Equation 15, together with the universal NA-independent expression for ζ Equation 13 form Equation 3 and describe the depth-dependent axial scaling when imaging under RIM.

### 2.5. Re-scaling microscopy data

When re-scaling microscopy data from the native *z*_*N*_ coordinate into the true depth coordinate *z* _*A*_, it is convenient to use the re-scaling factor as a function of the *nominal depth z*_*N*_ instead of the actual depth *z* _*A*_ utilized in Equation 13. In this case, a different dimensionless parameter *δ* can be used for the derivation:

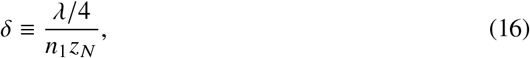

and Equation 12 can be re-written as:

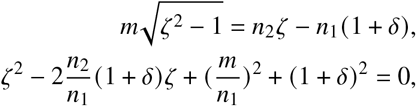

yielding the re-scaling factor as a function of *z*_*N*_ :

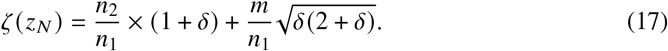

## 3. Methods

### 3.1. Wave-optics calculations

For the wave-optics calculations of the PSFs we adapted the most complete description from Hell *et al*. [17]. The radially symmetric fields (of both excitation and emission light) are represented as integrals over component Bessel-beams. The wave-optics is linked with geometrical optics through an elegant mathematical correspondence between Bessel beams and light-cone sections of the ray manifold [25, 26, 29]. The effects of RIM are taken into account as (RIM *dependent*) phase shifts of the component Bessel-beams, resulting in a new interference pattern of the field and thus in a new PSF shape. The (dimensionless) intensity of a focal spot can be found by computing three integrals *I*_0_, *I*_1_ and *I*_2_:

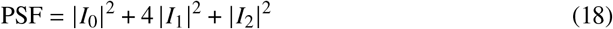

Integrals *I*_*i*_ each have a Bessel function of the first kind *J*_*i*_ (*r k*_*r*_ (*θ*)) weighted by some coefficients of geometric origin. The common factors are sin *θ* (from the Jacobian), 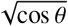 (from Abbe’s sine condition) and *e*^*i*Φ(*θ*)^ (the RIM induced phase shifts). The argument of the Bessel functions is *r k*_*r*_ (*θ*) = *r* × *n*_2_*k*_0_ sin *θ*. Taking into account the light polarization, the following factors appear: *I*_0_ contains a factor of (1 + cos *θ*), *I*_1_ a factor of sin *θ* and *I*_2_ a factor of 1 − cos *θ*. For each polarization orientation *s* and *p*, the partial transmission is taken into account as *t*_*s*_ (*θ*) and *t* _*p*_ (*θ*) coefficients. Thus, the polarization factors become: *t* _*p*_ (*θ*) + *t*_*s*_ (*θ*) cos *θ* in *I*_0_, *t*_*s*_ (*θ*) sin *θ* in *I*_1_ and *t* _*p*_ (*θ*) − *t*_*s*_ (*θ*) cos *θ* in *I*_2_. The integrals then become:

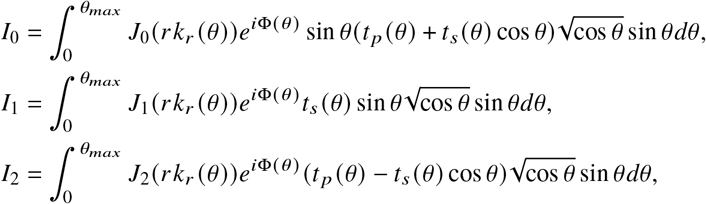

with *θ*_max_ = Re(arcsin NA/*n*_2_) the maximum angle of the light cone, and the phase function:

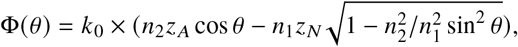

and the transmission coefficients *t*_*s*_ (*θ*) and *t* _*p*_ (*θ*):

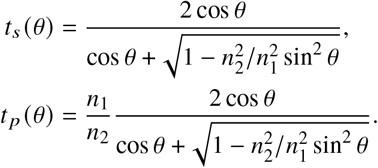

The computations of the integrals were performed through a MatLab script producing 2D PSF cross-sections (*r* × *z*_*N*_, 10 × 20 µm) on a *N*^2^ = 2048 × 2048 pixel grid from *M* = 2000 Bessel-beam components. To minimize the number of Bessel function invocations, each of the integrals *I*_*i*_ was represented as a double matrix product:

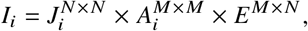

where 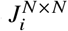 is the matrix containing the Bessel function, 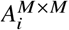 is the diagonal matrix created from all the position independent factors and *E* ^*M*×*N*^ is the matrix containing the *z*_*N*_ dependent phase change of *M* Bessel-beams.

For every RIM condition, a series of 2D PSF cross-sections (lateral × axial, XZ planes) was generated with the actual depth *z* _*A*_ changing from 0.2 to 5 µm with a step of 0.2 µm. Step sizes of 0.5 µm and 1.0 µm were used respectively from 5.5 to 25 µm and from 26 to 150 µm. The axial center of the calculated PSF frame was shifted with the actual depth *z* _*A*_ using the paraxial scaling factor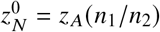. In the lateral center of each frame, the *Z* profile was extracted and passed through a Butterworth low-pass filter (5^th^ order, sampling and critical frequencies *f*_*s*_ = 0.1 nm^−1^, *f*_*c*_ = 0.003 nm^−1^) to remove high frequency ringing oscillations. The axial focal position 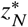 was determined by taking the maximum of the filtered profile and the re-scaling factor was computed as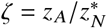.

### 3.2. Axial scaling measurements

We measured the depth-dependent axial scaling using a fluorescence microscope setup described earlier [9], for different refractive index mismatches and numerical apertures, as listed in Table 1. A schematic of the measurement scheme is shown in Figure 4a, where a sapphire ball (BA, Ceratec, 2 mm diameter, grade 10) is separated from a coverslip (CS, Thorlabs #CG15CH2, 22 × 22 mm, #1.5H thickness). The sapphire ball was glued to a glass strip (GS) with a small droplet of UV curable glue (Norland Optical Adhesive 63). Fluorescent beads (FB, diameter 190 nm, excitation 480 nm, emission 520 nm, Bangs Labs #FSDG002) were dropcasted on the coverslip using a micropipette. Both surfaces were cleaned with acetone, isopropyl alcohol and dried with nitrogen. The glued sapphire ball was dipped in the solution and dried three times to apply the beads to the surface. Beads on both surfaces were imaged by acquiring a z-stack with the fluorescence microscope using an excitation wavelength of 485 nm (Lumencor SPECTRA X), dichroic mirror (Semrock #FF410/504/582/669-Di01-25×36) and an emission filter (Semrock #FF01-525/30-25). The objective lens was mounted upright and was moved by stick-slip piezo positioners (SmarAct GmbH) in *X, Y* and *Z*. The correction collar setting of the OL was optimized to minimize spherical aberrations when imaging beads on the coverslip. The glass strip with sapphire ball was mounted to the sample shuttle holder (SSH), Figure 4b. This holder was moved by piezo positioners, and hence the distance between the coverslip and the ball was controlled. Details on the positioning systems can be found in [9].

**Table 1.** Overview of microscope objectives and refractive indices in the measurements.

| NA | Nikon # | $M$ | Pixel size [nm] | FOV [$\mu\text{m}$] | Correction collar | $n_1$ | $n_2$ |
| --- | --- | --- | --- | --- | --- | --- | --- |
| 0.70 | MRH08630 | $60\times$ | 108.3 | 110.9 | Yes | 1.0 | 1.0 |
|  |  |  |  |  |  | 1.0 | 1.34 |
|  |  |  |  |  |  | 1.0 | 1.52 |
| 0.85 | MUE35900 | $100\times$ | 65 | 133.12 | Yes | 1.0 | 1.0 |
|  |  |  |  |  |  | 1.0 | 1.34 |
|  |  |  |  |  |  | 1.0 | 1.52 |
| 0.95 | MRD00605 | $60\times$ | 108.3 | 110.9 | Yes | 1.0 | 1.0 |
|  |  |  |  |  |  | 1.0 | 1.34 |
|  |  |  |  |  |  | 1.0 | 1.52 |
| 1.25 | MRD77400 | $40\times$ | 162.5 | 166.4 | No | 1.34 | 1.34 |
| 1.40 | MRD01901 | $100\times$ | 65 | 133.12 | No | 1.52 | 1.52 |
|  |  |  |  |  |  | 1.52 | 1.34 |
|  |  |  |  |  |  | 1.52 | 1.0 |

**Fig. 4.**
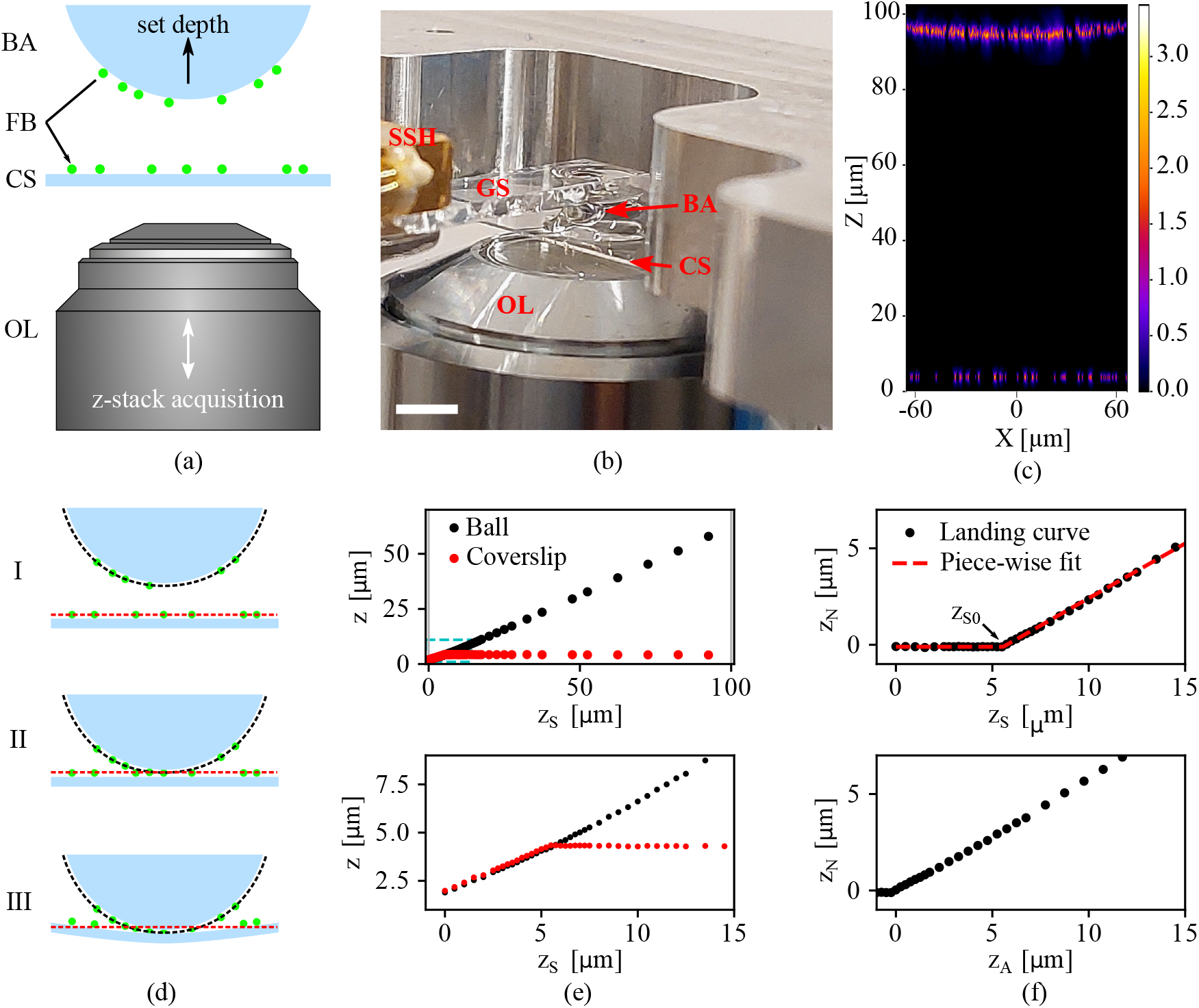
The experimental setup, and data analysis procedure, used to measure depth-dependent axial scaling. **(a)** Schematic showing the movable optical objective lens (OL) and sapphire ball (BA) at opposite sides of the static coverslip (CS), with fluorescent beads (FB) on their surfaces. **(b)** Photograph showing the objective lens (OL), coverslip (CS) and sapphire ball (BA). The ball is glued to a glass strip (GS), which is mounted to the sample shuttle holder (SSH). Scale bar 3 mm. **(c)** Maximum intensity projection along the *Y* axis of a recorded z-stack where *n*_1_ = *n*_2_ = 1.0. The bottom fluorescent signal originates from beads residing on the coverslip, and the signal at the top from beads on the sapphire ball, shown by the visible curvature. **(d)** Schematic showing three different distances between ball and coverslip, going from clear separation (I) to contact (II) and deformation of the coverslip (III). The dashed lines indicate the position of both ball (red) and coverslip (black). **(e)** The position of the coverslip and ball versus the set ball-coverslip distance *z*_*S*_ (top) and the same data near the coverslip (bottom). **(f)** The landing curve (in *z*_*N*_ coordinates), plotted against the set depth *z*_*S*_ (top). The fit values from a piecewise linear fit are used to determine the contact point between ball and coverslip, and this yields the observed focal depth *z*_*N*_ versus the actual depth *z* _*A*_ from which the re-scaling factor ζ = *z* _*A*_/*z*_*N*_ was calculated (bottom).

We used different objective lenses, but all were compatible with the Nikon CFI60 optical system, and were used together with a 200 mm tube lens (Nikon #MXA20696). Care was taken to center the pupil of the objective lens to the center of the tube lens, after which the objective lens was solely moved along the optical axis. With the distance between ball and coverslip set, a z-stack was acquired by moving the objective lens. In Figure 4c the maximum intensity projection along *Y* is shown for such a z-stack, where the fluorescent beads were imaged using a 100 ×, 0.85 NA objective (Nikon #MUE35900) in the absence of a refractive index mismatch (*n*_1_ = *n*_2_ = 1.0). The bottom fluorescent signal originates from fluorescent beads residing on the coverslip. The signal at the top comes from the beads on the sapphire ball, as recognized by the visible curvature.

The AFP was varied by changing the distance between ball and coverslip, and the contact point between them was first approximately set by having both the fluorescent beads on the coverslip and bottom side of the ball in focus of the microscope. Next, the starting AFP (*z* _*A*_, typically around 100 µm) was set by moving the ball away from the coverslip and the AFP was reduced step by step (see Figure 4d, I through III). At each set depth *z* _*A*_, a z-stack is recorded around the coverslip and ball separately (by moving the objective lens, − 4 to 8 µm and − 8 to 8 µm respectively, both with a step size of 0.25 µm). To ensure the second z-stack was acquired around the beads on the surface of the ball, we used ζ_marginal_ and ζ_paraxial_ estimates for < 20 µm and, > 20 µm respectively. At the end of the measurement sequence the ball was forced into contact with the coverslip, slightly deforming it by setting negative values of AFP on purpose (to about − 2 µm), see Figure 4d, III.

The 3D positions of all fluorescent beads in the recorded z-stack were acquired using the PSF Extractor software [9, 30] and the *X, Y* and *Z* locations of all fluorescent beads were extracted. From these, the position of the coverslip *z*_*CS*_ was determined by taking the median of the *z* positions of the beads on the cover slip and a least squares paraboloid fit on the coordinates of the beads residing on the ball surface was used to determine the lowest point of the ball *z*_*B*_. These *z* positions are plotted against the set depth *z*_*S*_ in Figure 4e for the ball (black) and coverslip (red). Around 5.5 µm the ball and coverslip make contact, effectively deforming the coverslip at lower set depths. The coverslip position was subtracted from the lowest point of the ball (*z*_*N*_ = *z*_*B*_ − *z*_*CS*_) and plotted against (*z*_*S*_), see Figure 4f (top). A piece wise linear function of the form

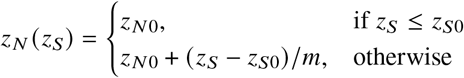

is used to determine the offset *z*_*N*0_ between coverslip and ball, the inflection point *z*_*S*0_ and the slope of the curve *m*. The fit is performed using a Levenberg-Marquardt non-linear least squares algorithm. We fit the data from − 2 to 20 µm around the inflection point, and pass the measured values to the uncertainty parameter *sigma* in scipy.optimize.curve_fit [31]. This artificially increases the uncertainty with increasing depth, and ensures that the inflection point can be found reproducibly for all datasets. The fit values are used to calculate the AFP for each set depth *z*_*A*_ = *z*_*S*_ − *z*_*N*0_ + *mz*_*S*0_. The result is shown in Figure 4f (bottom), where the observed nominal depth *z*_*N*_ versus the actual depth *z* _*A*_ is plotted, from which the re-scaling factor (ζ = *z* _*A*_ /*z*_*N*_) was computed.

### 3.3. Re-scaling of 3D microscopy data

To demonstrate and test the re-scaling of 3D microscopy data, we recorded z-stacks of fluorescent beads embedded in an agarose hydrogel for *n*_1_ < *n*_2_, *n*_1_ = *n*_2_ and *n*_1_ > *n*_2_, where the *n*_1_ = *n*_2_ case serves as a ground truth for the re-scaling of the mismatched cases. To prepare the sample, we first coated a cover slip with fluorescent beads (190 nm, excitation 480 nm, emission 520 nm, Bangs Labs #FSDG002) by dropcasting from isopropyl alcohol (IPA). Next, an agarose gel was prepared (2w%) by mixing Milli-Q water and agarose (Low EEO, Fisher Scientific) and heating to boil in a microwave. 25 µL of the same beads in IPA were added to 225 mL agarose gel while warm. The mixture was pipetted into a glass sample cell composed of a microscope slide to which two small pieces of microscope slide as spacers were glued (using Norland 81 optical adhesive) roughly 0.5 cm apart, on top of which a #1.5 cover slip (180 µm thickness) was glued. After filling the cell with the agarose gel containing the beads, the cell was sealed with optical adhesive. After each gluing step, the sample was cured by exposure to UV light (∼ 350 nm) for 90 s, and during the last curing step the beads were protected using a piece of aluminum foil.

Confocal z-stacks of the same volume were repeatedly recorded with different objectives using a Nikon C2-SHS C2si confocal on a Nikon Eclipse Ti inverted microscope with a 488 nm excitation laser. If the microscope objective had a correction collar, we optimized its position to minimize spherical aberrations while inspecting beads on the cover slip before the recording of the z-stack. We used the following procedure to record the z-stacks (with a 500 nm z step size). On the back of the sample (i.e. the side of the microscope slide) a spot was drawn using a felt marker. We then imaged this spot in bright field using a 10 × /0.4 NA air objective and chose a reference feature in this spot to find back the same position in the sample. Next, we switched to the 100 × /1.4 NA oil objective, and recorded a confocal z-stack including the cover slip interface. We then removed the immersion liquid from the sample, and found the same position again using the 10 × /0.4 air objective. At this position, we again recorded a z-stack, but now with the 100 × /0.85 NA air objective. Finally, we recorded at the same position a (ground truth) z-stack using a 40 × /1.25 NA water objective. For single bead comparison, we cropped volumes in the three stacks comprising the same beads in the ground truth (water immersion) stack and the axially deformed stacks (air, oil immersion). We determined the cover-slip-to-sample interface as the intensity peak of the beads deposited on the cover slip and cropped the z-stack above the cover slip.

Re-scaling of the data was done either using the linear theories [4, 13, 15, 16] where the voxel size in *z* was simply corrected, or using the depth-dependent re-scaling. For the latter, the AFP of each slide in the stack was calculated using equation 17, with the Lyakin scaling factor as the critical value instead of equation 15. Next, a linear z-stack was generated with a z step size corresponding to the original stack and a range corresponding to the final AFP. The intensities in this z-stack were interpolated (using inverse distance weighting) from the two nearest slices in the (non-linear) AFP z-stack. A Jupyter Notebook to perform the linear and non-linear re-scaling is available at [32].

### 3.4. Refractive index measurements

The refractive indices of the immersion oils used in this study were measured using an Abbe refractometer (Atago 3T). The indices were measured at *T* = 25 °C and at a wavelength *λ*_*D*_ = 589.3 nm. Using a measured dispersion value, the indices were converted using Cauchy’s relation to *λ* = 520 nm [33]. This resulted in the following refractive indices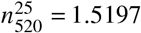 (Type DF immersion oil, Cargille) and 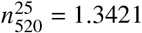 (Immersol™ W 2010 immersion oil, Zeiss).

## 4. Results

### 4.1. Validation through wave-optics calculations

We first validate the derived analytical expression, by comparing it to wave-optics calculations, performed as described in subsection 3.1. This is shown in Figure 5a and b respectively for RIM contrasts of *n*_1_ = 1.0 → *n*_2_ = 1.33 and *n*_1_ = 1.52 → *n*_2_ = 1.33. The axial re-scaling factor ζ is plotted versus depth, as obtained through analytical expression Equation 3 (colored solid) and as calculated using wave-optics (colored dashed).

**Fig. 5.**
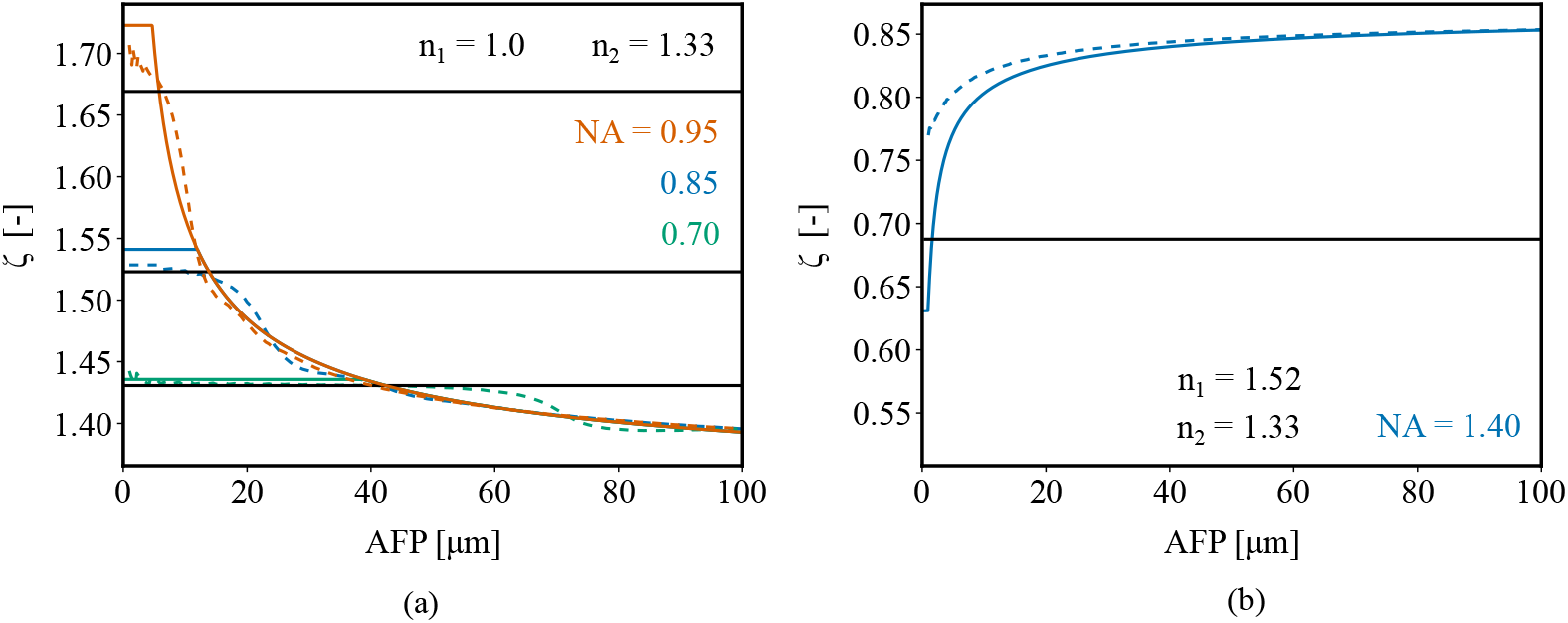
Axial re-scaling factor ζ versus AFP for different NAs and RIM contrasts, obtained through analytical expression Equation 3 (colored solid) and as calculated using wave-optics (colored dashed). For each NA the re-scaling factor as calculated by Lyakin *et al*. is included as a black solid line [16]. **(a)** ζ as a function of AFP for NAs of 0.95, 0.85 and 0.70 at *n*_1_ = 1.0 and *n*_2_ = 1.33. **(b)** ζ as a function of AFP for an NA of 1.4 at *n*_1_ = 1.52 and *n*_2_= 1.33.

For *n*_1_ < *n*_2_ (Figure 5a) the re-scaling factor increases at shallow depths, where the re-scaling (constant) maximum is determined by the numerical aperture of the optical system (i.e. the *z* _*A*_-independent value of ζ_crit_). At increasing depths it levels towards the *n*_2_ / *n*_1_ limit, following the universal (i.e. NA-independent) curve ζ_univ_. The wave-optics calculations deviate from the analytical expression due to some oscillations remaining after data processing of the computed 2D PSFs. Despite this, they follow the general trend in the NA-independent regime (ζ_univ_ in Equation 3) for the different NAs plotted. At shallow depths, the plateau from the analytical expression (ζ_crit_ in Equation 3) is reproduced by the wave-optics calculations, though the exact re-scaling factor value fits between the analytical expression and the re-scaling factor as calculated by Lyakin *et al*. [16].

In the case of *n*_1_ > *n*_2_ (Figure 5b) the re-scaling factor as given by Equation 3, decreases at shallow depths, also reaching a plateau value determined by the numerical aperture. For this RIM contrast the plateau is very small, yielding a fast changing re-scaling factor in the first 10 µm after the refractive index interface. Imaging deeper, the re-scaling factor again levels towards the *n*_2_ / *n*_1_ limit. The wave-optics calculations show less extreme axial scaling at shallow depths, while we find good agreement for depths > 15 µm.

### 4.2. Validation through experiments

Five different objective lenses are used to measure depth-dependent axial re-scaling factors as explained in subsection 3.2, varying in numerical aperture, immersion refractive index *n*_1_ and magnification. Immersol^*T M*^ W 2010 immersion oil (Zeiss,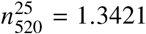) and Type DF immersion oil (Cargille,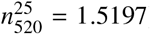) were used in experiments. We will first discuss two cases in detail below, *n*_1_ = 1.0 → *n*_2_ = 1.34 for a NA of 0.85 and *n*_1_ = 1.52 → *n*_2_ = 1.34 for a NA of 1.4. For the other cases listed in Table 1, figures are presented in Supplement 1 and will be discussed afterwards.

#### 4.2.1. NA=0.85, *n*_1_ = 1.0, *n*_2_ = 1.34

In Figure 6, the axial re-scaling factor ζ is plotted against the actual depth *z* _*A*_. The measurements (solid blue dots) are plotted alongside the wave-optics calculations (dashed blue lines), analytical solution (solid black lines) and literature theories (solid colored lines). From our individual sets of measurement data, the mean (solid black dots) is computed by binning along *z* _*A*_, plotted in the center of each bin (bin sizes of 1 and 10 µm respectively for 0 to 10 µm and 10 to 100 µm). We estimate an upper error limit of (100 nm /*z*_*N*_ + 100 nm / *z*_*A*_) ζ, in determining the various *z* positions, composed of errors from: (i) positioning of the piezo stages, (ii) determining the position of ball and coverslip by fluorescent bead localization, and (iii) fitting uncertainties in the landing curve. The total measured error for the re-scaling factor (shaded blue area) is the sum of the rolling standard deviation of the measurements and the upper error limit. It increases significantly at shallow depths, which is confirmed by acquiring data without RIM present, and applying the same data analysis as shown in Figures S4 and S10.

**Fig. 6.**
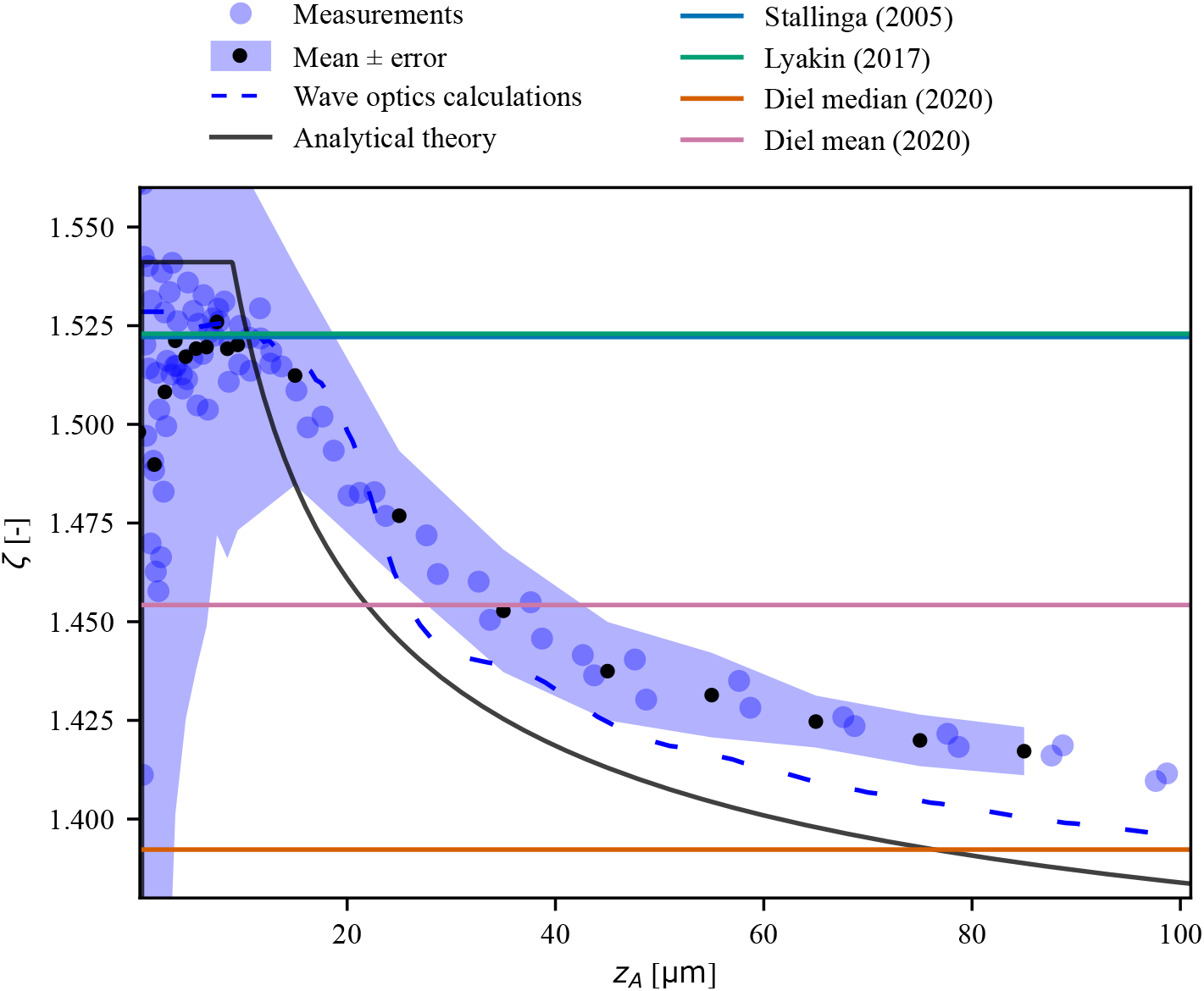
Axial re-scaling factor ζ versus AFP for an NA of 0.85, imaging under a RIM contrast of *n*_1_ = 1.0 → *n*_2_ = 1.34. Measurement data (solid blue dots) are plotted alongside the wave-optics calculations (dashed blue lines), analytical solution (solid black lines, Equation 3) and depth-independent theories (solid lines). Two sets of measurement data are plotted individually and from this data the mean (solid black dots) is computed by binning along *z* _*A*_, plotted in the center of each bin (bin sizes of 1 and 10 µm respectively for 0 to 10 µm and 10 to 100 µm). The measurement error (shaded blue area) is the sum of the measurements standard deviation and an estimated upper error limit.

Results are shown for a RIM contrast of *n*_1_ = 1.0 → *n*_2_ = 1.34 in Figure 6, where the uncertainty in the measurement data increases at shallow depths, while at larger depths the measurements capture the decay of the re-scaling factor but are slightly higher than the wave-optics calculations and analytical theory. Although we have taken care to align the optical system, we do expect some misalignment and/or residual optical aberrations to be the cause of this deviation. Especially, residual spherical aberration induced by a non-perfect correction collar setting could influence the measurements.

Looking at the exact value of the plateau at shallow depths, the analytical expression overestimates the re-scaling factor compared with the measurements and wave-optics calculations. In fact, both re-scaling factors from Lyakin and Stallinga give better agreement with the measured data [15, 16] and in practical terms seems to provide a better value for ζ_crit_. At extreme depths of *z* _*A*_ = 10 mm the depth-dependent re-scaling factor approaches the *n*_2_ / *n*_1_ = 1.34 limit set by Carlsson *et al*. (not shown in Figure 6) [13]. At more realistic imaging depths of 100 µm, the median from Diel coincides with the depth-dependent re-scaling factor and their mean gives a good agreement at *z* _*A*_ = 40 µm.

#### 4.2.2. NA=1.4, *n*_1_ = 1.52, *n*_2_ = 1.34

In Figure 7, the axial re-scaling factor ζ is plotted against the actual depth *z* _*A*_. We find good agreement between measurement data and the analytical theory when imaging under a RIM contrast of *n*_1_ = 1.52 → *n*_2_ = 1.34, as show in Figure 7. The measurements presented agree with the analytical theory and only start to drop more at shallow depths (< 4 µm). The relative error when approaching *z* _*A*_ = 0 increases for *z* _*A*_ and *z*_*N*_ as the error in both is affected by determining the point at which the sapphire ball touches the coverslip. We have also included re-scaling factors as measured by Petrov *et al*. where they use a method with a much smaller measurement errors (see also inset) [23]. Combining the measurement data from Petrov at shallow depths, and our own measurements data at larger depths, we find very good agreement with the analytical theory presented. Although the analytical expression provides a plateau for the re-scaling factor ζ_crit_ = 0.64, this is not reproduced in the measurement data from Petrov.

**Fig. 7.**
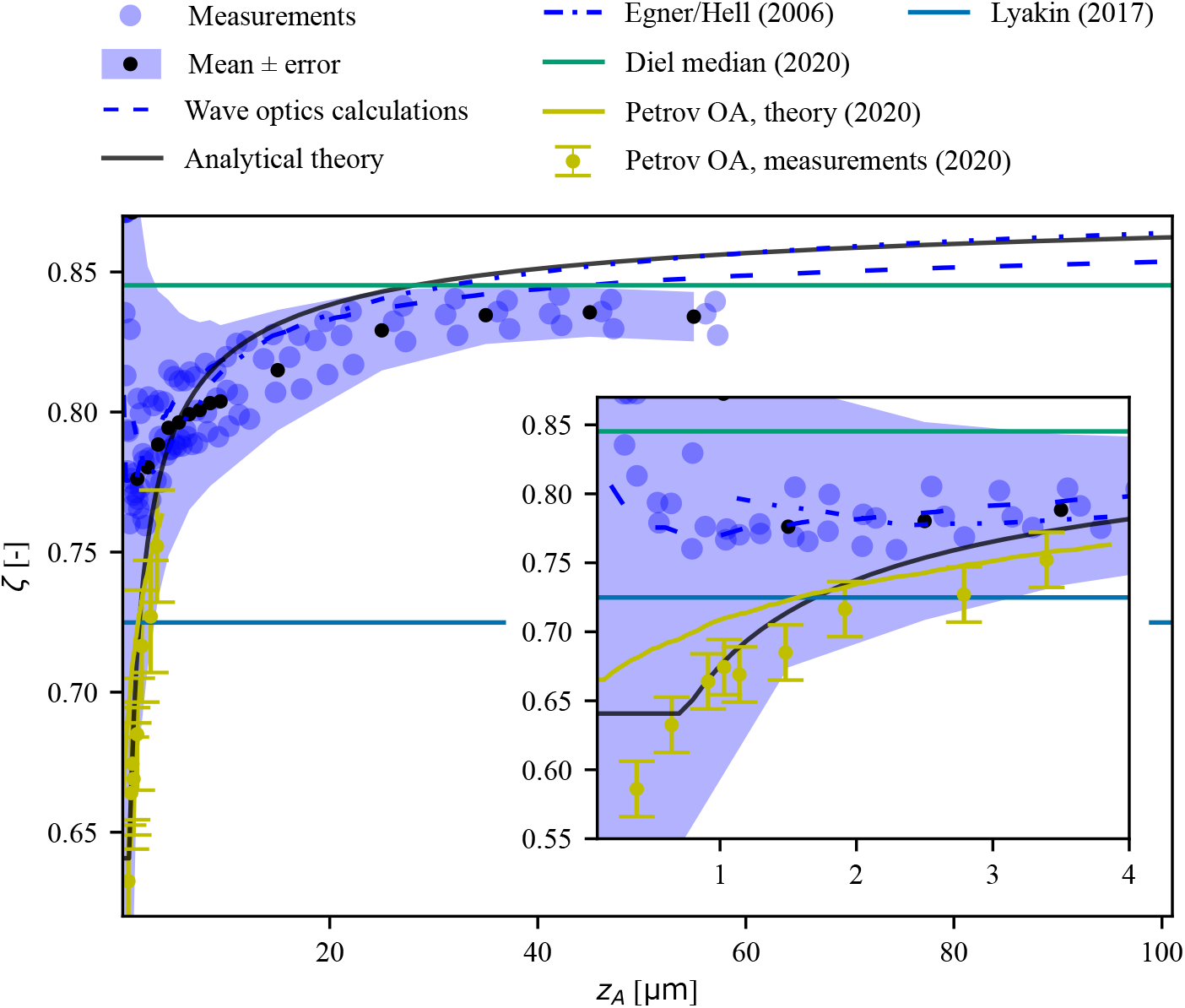
Axial re-scaling factor ζ versus AFP for an NA of 1.4, imaging under a RIM contrast of *n*_1_ = 1.52 → *n*_2_ = 1.34. Measurement data (solid blue dots) are plotted alongside the wave-optics calculations (dashed blue lines), analytical solution (solid black lines, Equation 3) and depth-independent theories (solid lines). Three sets of measurement data are plotted individually and from this data the mean (solid black dots) is computed by binning along *z* _*A*_, plotted in the center of each bin (bin sizes of 1 and 10 µm respectively for 0 to 10 µm and 10 to 100 µm). The measurement error (shaded blue area) is the sum of the measurements standard deviation and an estimated upper error limit. Measured data and theory as presented by Petrov *et al*. and wave-optics calculations from Egner & Hell are included [19, 23].

Our wave-optics calculations also match the analytical theory at larger depths, but deviate below *z* _*A*_ = 4 µm. At the same time, they do overlap with wave-optics calculations from Egner & Hell [19]. At these shallow depths, an offset of a few tens of nanometers in the NFP has a dramatic effect on the re-scaling factor and hence super-critical angle fluorescence (SAF) effects are significant [34]. Although SAF is not included in the analytical derivation, it is considered in the wave-optics calculations through the Fresnel transmission coefficients when using the vector PSF model. The axial deformation is, however, enhanced by SAF, as the NFP shifts <40 nm away from the objective [34]. Therefore, the difference in the re-scaling factor between the wave-optics calculations and the analytical theory has already been reduced by the incorporation of SAF in the calculations.

Up to depths of *z* _*A*_ = 1 µm, there is a ∼ 250 nm difference in the NFP found between ζ = 0.64 (analytical theory) and ζ = 0.78 (wave-optics calculations). This is too large to be caused by the chosen method of determining the focal position (i.e. maximum axial intensity [19] vs. minimal lateral width [23]). As such, we do not have a fitting explanation for the discrepancy between the wave-optics calculations and the analytical theory along with measurements from Petrov *et al*..

#### 4.2.3. Other imaging scenarios

The measurement data for the remaining cases listed in Table 1, are shown in Figures S1 through S11, where the re-scaling factor ζ is plotted against *z* _*A*_, alongside the wave-optics calculations, analytical solution and depth-independent theories. For each objective lens used we acquired data without a refractive index mismatch present and applied the same data analysis procedures as for axially scaled measurements (Figures S1, S4, S6, S9, S10). In all cases, the spread in the measurement data increases below *z* _*A*_ = 10 µm as uncertainties in determining both *z* _*A*_ and *z*_*N*_ increase. At higher depths, no reproducible deviations away from ζ = 1.0 are found in these data, and we note a maximum error of about 2 % in the measurements. The data shown in Figure S9 is of special interest where we have used a water immersion objective to verify our measurement method and analysis when having immersion oil (*n* = 1.34) present between the coverslip and sapphire ball. We find a mean re-scaling factor of 0.99 ± 0.05 (standard deviation) over the entire depth range, and apart from the increased scatter near *z* _*A*_ = 0, the re-scaling factor remains approximately constant (ζ = 1.0) as a function of depth for each individual dataset.

The objective lens with the lowest numerical aperture which we used in our measurements had a NA of 0.7, of which the results are presented in Figure S2 and S3 (*n*_1_ = 1.0 → *n*_2_ = 1.34 & *n*_1_ = 1.0 → *n*_2_ = 1.52). In both cases, the wave-optics calculations reproduce the analytical expression, while the measurements only reproduce the general trend for higher RIM contrast of *n*_1_ = 1.0 → *n*_2_ = 1.52. In the case of *n*_1_ = 1.0 → *n*_2_ = 1.34, the measurement uncertainties obscure a clear trend in the measurement data.

The RIM of *n*_1_ = 1.0 → *n*_2_ = 1.52 for the 0.85 NA objective lens is shown in Figure S5. Both the wave-optics calculations and measurement data reproduce the behavior of the re-scaling factor as predicted by the analytical expression. Comparing against the measurement data, Lyakin and Stallinga again provide the best critical value ζ_crit_.

In the case of the 0.95 NA air objective (Figures S7 and S8), again the wave-optics calculations reproduce the analytical expression over the entire depth range. For the measured data however, the trend of the NA independent regime ζ_universal_ is reproduced, but the measurements fail to replicate the strong increase to the plateau given by ζ_crit_ below *z* _*A*_ = 20 µm. Along with depth-dependent re-scaling being more prominently present with increasing NA, so is the sensitivity to any remaining (i.e. spherical or tilt) aberrations in the optical setup. We have taken care to minimize these, but we estimate that residual aberrations lower the effective NA, obscuring the rise to the plateau in the measurements.

For the 1.4 NA oil immersion objective, we present the RIM case of *n*_1_ = 1.52 → *n*_2_ = 1.0 in Figure S11. Again the general depth-dependent behavior is reproduced in the measurements, but the re-scaling factor is underestimated at the coverslip while it is overestimated when imaging deeper into the sample.

### 4.3. Axial re-scaling of 3D microscopy data

To test axial re-scaling of 3D microscopy data using our depth-dependent theory, we recorded confocal z-stacks of beads suspended in an agarose hydrogel, from the cover slip to 100 µm depth (*z*_*N*_). These z-stacks were recorded using the 100 × /0.85 NA air, 40 × /1.25 NA water and 100 × /1.4 oil objectives (see Table 1). As the refractive index of the agarose gel (*n*_2_ = 1.3356 [35]) is very close to the refractive index of the immersion medium of the water objective, we can use these z-stacks as a ground truth for the axial re-scaling of the data. We recorded the z-stacks at the exact same location in the sample to allow for direct comparison of the same beads in different imaging scenarios.

#### 4.3.1. *n*_1_ < *n*_2_

The case *n*_1_ < *n*_2_ is particularly relevant for cryo-fluorescence microscopy where air immersion is used to observe a frozen or vitrified sample. Thus, we evaluate a typical situation with NA = 0.85, *n*_1_ = 1.0 and *n*_2_ = 1.336 [9]. Before re-scaling the data, we quantitatively compare the re-scaling factors of the depth-dependent and linear theories, as plotted in Figure S12. As we have found that the theories of Lyakin and Stallinga result in approximately equal re-scaling factors, we choose to plot only the Lyakin re-scaling factor. As the critical value of the depth-dependent re-scaling factor, we use the Lyakin re-scaling factor. Therefore, the depth-dependent re-scaling factor is 1.532 close to the cover slip and falls off to ∼ 1.4 deeper into the sample (see Figure S12a). When plotting the focal shift Δ *f* as a function of depth (AFP) we see that depth-dependent focal shift equals the Lyakin focal shift close to the cover slip and gradually transitions to the Diel (median) focal shift at larger depth (see Figure S12b).

To see how significant the difference between the theories is for this imaging scenario, we plot the absolute and relative difference between the depth-dependent and linear theories as function of depth (AFP). Figure S12c shows that for Lyakin the difference is obviously zero near the cover slip, but quickly increases to a large difference (>1 µm) after a depth of >25 µm. On the other hand, Diel (median) differs significantly (∼ 1 µm) at a depth of 20 µm, but is the linear theory with the smallest relative difference at >50 µm depth. Finally, Diel (mean) is in between the other two linear theories, having a smaller difference close to the cover slip than Diel (median), and a smaller difference at large depth than Lyakin. Where the relative difference between the depth-dependent theory and the theories of Lyakin and Diel median can exceed 5 %, the relative difference for the Diel mean theory is below 3 % (see Figure S12d).

Figure 8 shows the re-scaling of the 3D microscopy data of beads in an agarose hydrogel with NA = 0.85, *n*_1_ = 1.0 and *n*_2_ = 1.336. Overlays are plotted of the re-scaled z-stacks (grays) and the ground truth z-stack recorded with the water immersion objective (reds). As *n*_1_ < *n*_2_, the recorded z-stack was stretched in the axial direction to return to the ground truth axial distances. Deep into the sample (*z*_*A*_ ≈ 90 µm), Diel (mean) shows the largest re-scaling error where the z-stack is overstretched, whereas the depth-dependent and Diel (median) re-scaling collapses onto the ground truth data, although the PSF is elongated in the axial direction due to spherical aberrations induced by the refractive index mismatch. Closer to the cover slip (*z*_*A*_ ≈ 20 µm), both the depth-dependent and Diel (mean) re-scaling overlap with the ground truth z-stack, whereas Diel (median) has a slight (absolute) error as the axial distances have not been stretched enough. This agrees with the re-scaling factors plotted in Figure S12.

**Fig. 8.**
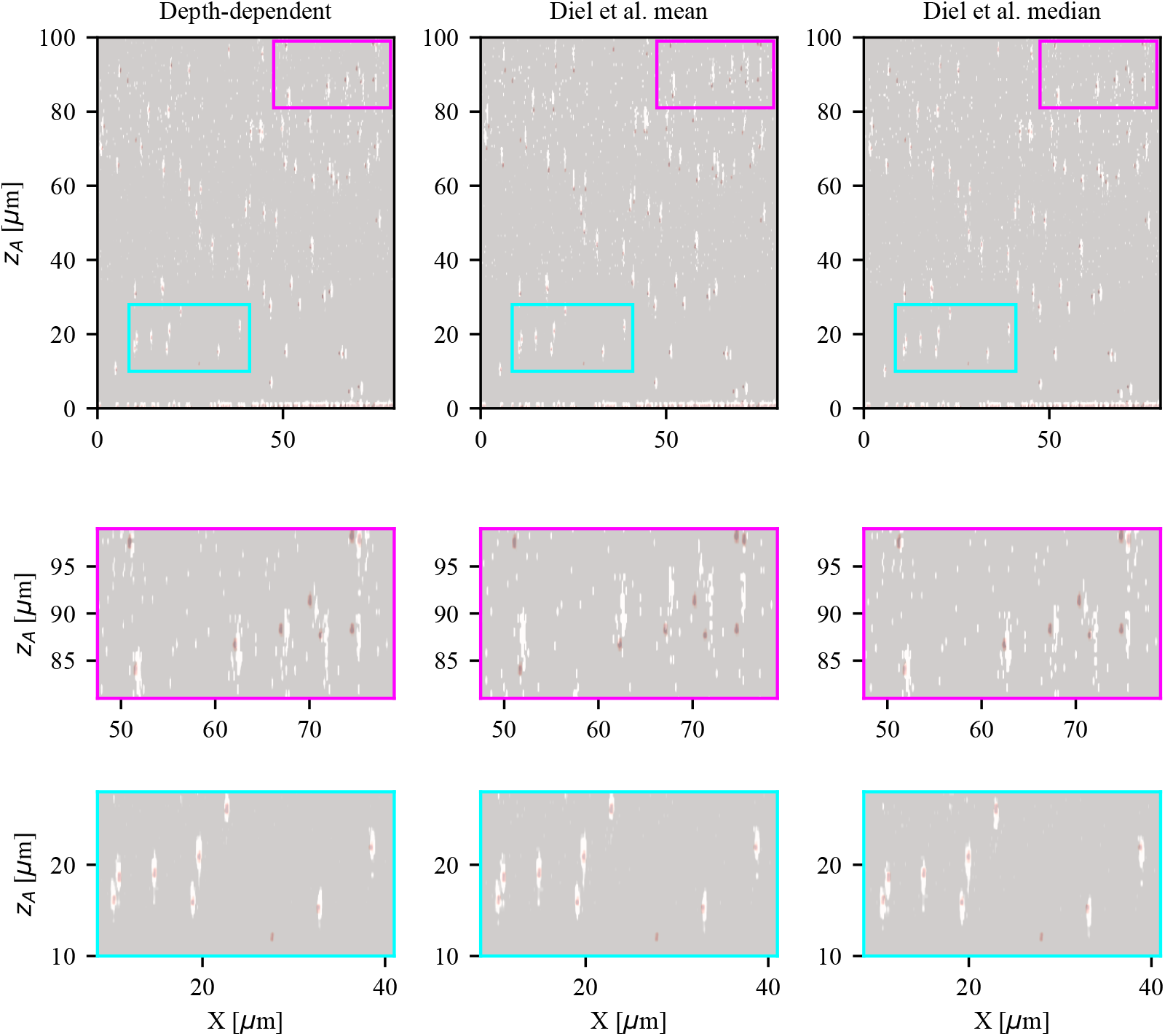
Axial re-scaling of 3D microscopy data with NA = 0.85, *n*_1_ = 1.0, *n*_2_ = 1.336 using the depth-dependent and linear theories. Maximum intensity projections along *Y* of the (re-scaled) confocal z-stack of beads embedded in an agarose hydrogel. The stacks have been re-scaled using the depth-dependent (left), Diel (mean) (middle) and Diel (median) (right) re-scaling factors. Overlays are plotted of the re-scaled z-stacks (grays) and the ground truth z-stack recorded with the water immersion objective (reds). The two bottom rows are cut-outs of the upper row, where the cut-outs at larger depths appear more noisy due to the intensity re-scaling due to fluorescence intensity loss at large depths. The beads are slightly displaced in *X* due to imperfections in the manual overlay of the z-stacks.

#### 4.3.2. *n*_1_ > *n*_2_

The case of *n*_1_ > *n*_2_ is relevant for oil immersion observation of samples in water. Thus, we here evaluate a typical situation with NA = 1.4, *n*_1_ = 1.52 and *n*_2_ = 1.336. We quantitatively compare the re-scaling factors of the depth-dependent and linear theories in Figure S13. As NA > *n*_2_, we cannot use Diel (mean). The depth-dependent re-scaling factor equals the Lyakin re-scaling factor close to the cover slip, but surpasses the Diel (median) at *z* _*A*_ ≈ 30 µm. The same trend is seen in the focal shift Δ *f*. In this imaging scenario, the axial positional difference between Lyakin and the depth-dependent re-scaling theory increases after a few micrometers in depth, resulting in a significant difference >6 %. While the absolute difference between Diel (median) and the depth-dependent theory is small close to the cover slip (<1 µm) and ∼ 2 µm um at large depths, the relative error is large (∼ 20 %) close to the cover slip and small at large depths (∼ 2 %).

In Figure 9 the re-scaling of the 3D microscopy data is plotted. As *n*_1_ > *n*_2_, the recorded z-stack was compressed in the axial direction to return to the ground truth axial distances. At larger depth (*z*_*A*_ ≈ 65 µm), the depth-dependent re-scaling nicely coincides with the ground truth, whereas the axial distances in the Diel (median) re-scaling has been compressed too much (see second row). Closer to the cover slip (*z*_*A*_ ≈ 15 µm) the depth-dependent re-scaling again nicely coincides with the ground truth, but the Diel (median) re-scaling has been compressed too little.

**Fig. 9.**
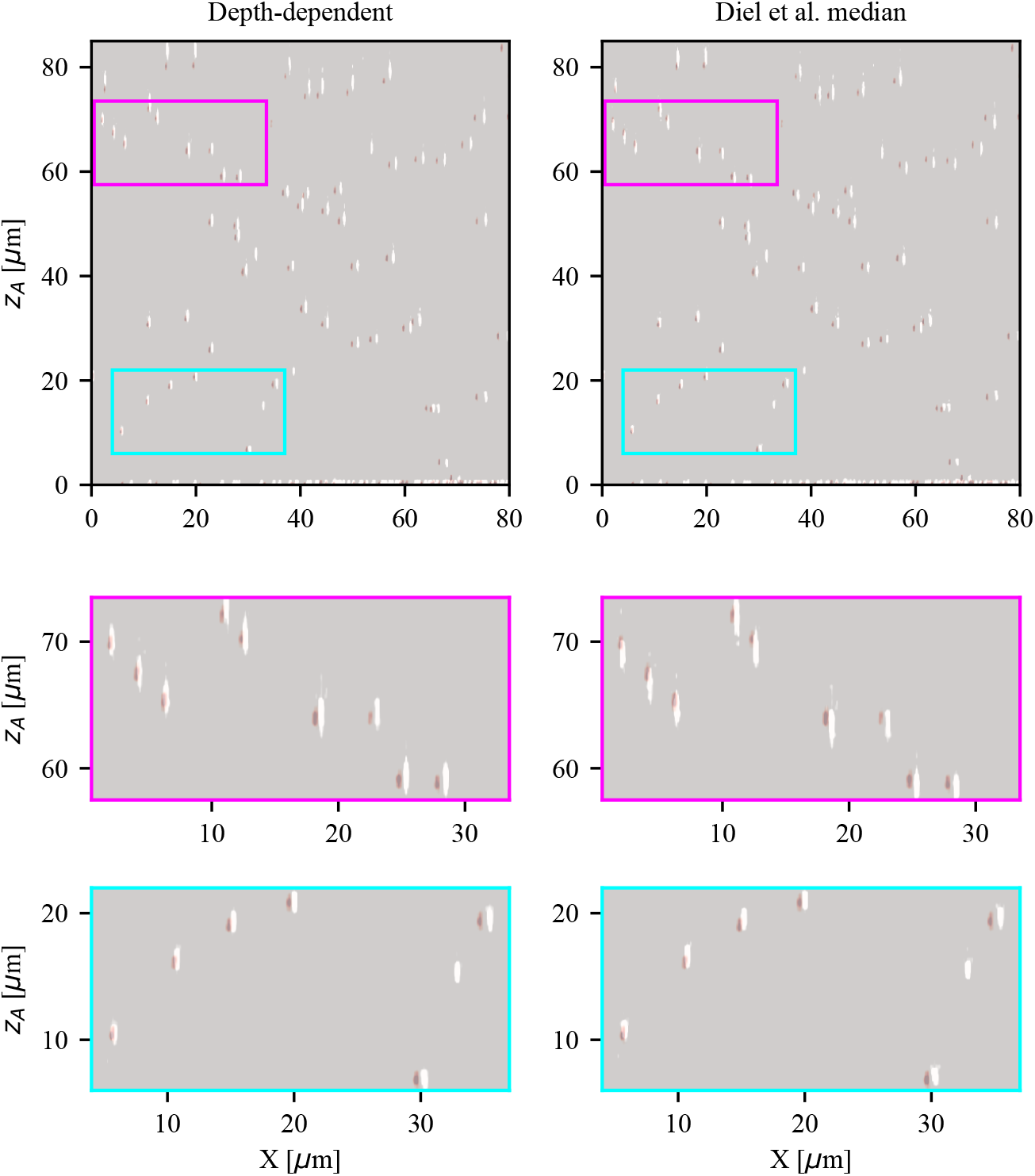
Axial re-scaling of 3D microscopy data with NA = 1.4, *n*_1_ = 1.52, *n*_2_ = 1.336. Maximum intensity projections along *Y* of the (re-scaled) confocal z-stack of beads embedded in an agarose hydrogel. The stacks have been re-scaled using the depth-dependent (left) and Diel (median) (right) re-scaling factors. Overlays are plotted of the re-scaled z-stacks (grays) and the ground truth z-stack recorded with the water immersion objective (reds). The two bottom rows are cut-outs of the upper row. The beads are slightly displaced in *X* due to imperfections in the manual overlay of the z-stacks.

#### 4.3.3. Quantitative comparison

To quantify how well the acquired data can be corrected using the depth-dependant re-scaling factor, we localize all individual fluorescent beads present in the data recorded using the PSF-Extractor software [30]. We compare the *z* position found in the re-scaled data to the ground truth value *z* _*A*_ and divide this difference over *z* _*A*_. This is shown in Figure 10 for both imaging scenarios, where we bin the data (bin size 10 µm), plot the mean of each bin along with the standard deviation as error bar, and plot the data centered with respect to the bin. In Figure 10a (NA = 0.85, *n*_1_ = 1.0, *n*_2_ = 1.336), we omit the first bin (0 to 10 µm) as it only contains 3 data points. The depth-dependent theory outperforms both Diel theories in the axial re-scaling of this data, where we note a maximum error of 2 % when using depth-dependent axial re-scaling, whereas the errors for the linear theories approaches 5 %.

**Fig. 10.**
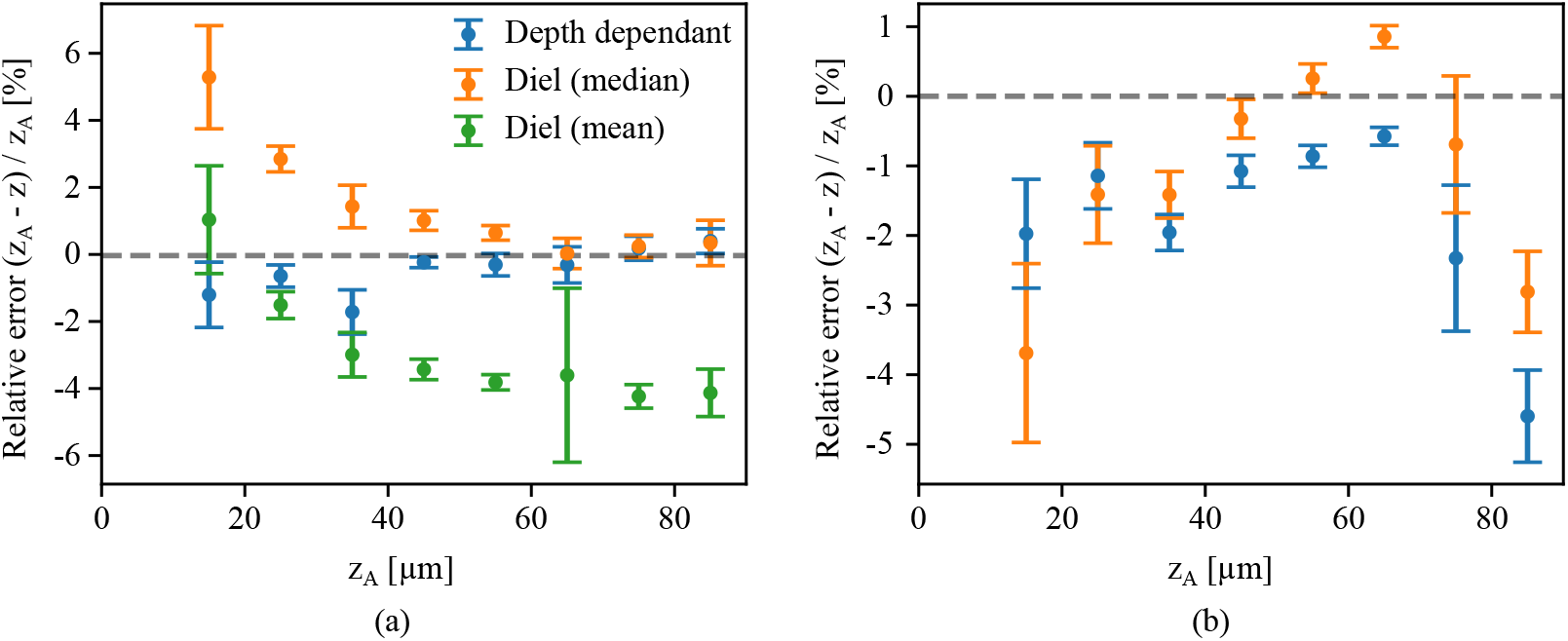
Relative error of the re-scaled *z* position compared to the ground truth *z* _*A*_ as recorded using a water immersion objective. The relative error is binned along *z* _*A*_ in bins of 10 µm and the mean of each bin is plotted in the center of each bin. The standard deviation is plotted as an error bar. The dashed gray line indicates 0 and acts as a guide to the eye. **(a)** Results for NA = 0.85, *n*_1_ = 1.0, *n*_2_ = 1.336, where we measure an error in the axial re-scaling using the depth-dependent theory below 2 %, whereas the linear theories result in errors close to 5 %. **(b)** Results for NA = 1.4, *n*_1_ = 1.52, *n*_2_ = 1.336, where we we find that the maximum error in the axial re-scaling using the depth-dependent and Diel (median) linear theory is 5 %.

The same analysis and data processing is done for NA = 1.4, *n*_1_ = 1.52, *n*_2_ = 1.336, to quantify the error in re-scaling using the Diel (median) and depth-dependent theory, see Figure 10b. There is not one theory that outperforms the others over the depths measured here. Still, the depth-dependent theory results in a smaller error closer to the cover slip, as is to be expected following Figure 7. In addition, this measurement did not include beads near the cover glass (*z* _*A*_ < 20 µm) where a relative difference up to 20 % between the two theories is expected (see Figure S13).

We should note that the objective lens used in this experiment did not have a correction collar. This means that we could not completely get rid of spherical aberration (SA) when imaging close to the cover slip. The presence of SA near the cover slip contradicts the optical conditions in the derivation of the analytical theory and therefore affects the legitimacy of the depth-dependent re-scaling factor, resulting in a higher re-scaling error. This shows that proper re-scaling also requires an optimization of the correction collar before the acquisition of the 3D microscopy data.

### 4.4. Software for depth-dependent re-scaling

#### 4.4.1. Re-scaling acquired z-stacks using Python

To re-scale z-stack acquired under RIM, we have written Python software which is available at https://github.com/hoogenboom-group/SF [32]. Jupyter Notebooks are used to read image data and, based on the specific imaging conditions, use inverse distance weighting to rescale the intensities along the axial coordinate correctly. The same axial pixel size from the original data is used, but depending on the RIM mismatch, the re-scaled z-stack will map a shorter or larger axial range.

#### 4.4.2. Interactive online tool for plotting the re-scaling factor versus imaging depth

We have made an online interactive tool where the depth-dependent re-scaling factor can be plotted and compared to existing depth independent scaling theories using Plotly Dash [36], the source code of which is available at [32]. The code is accessible via the URL https://axialscaling.pythonanywhere.com/. The refractive indices *n*_1_ and *n*_2_, the NA and the wavelength *λ* can be varied. In addition, the focal shift Δ *f* = can be plotted for the depth-dependent re-scaling factor, as well as for two depth independent scaling theories. Both the re-scaling factors and the focal shift resulting from the depth-dependent theory can be exported to file. All the measurement data and wave optics calculation results found in this manuscript and the accompanying supplemental materials are also included in the interactive plot.

## 5. Discussion

The various re-scaling theories found in the literature can be understood as a result of different (overt or covert) assumptions on which maximum constructive interference contribution is leading. For example, Lyakin *et al*. used an analysis similar to ours, but explicitly sets the critical point *θ*^*∗*^ such that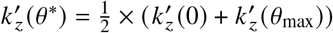, and then from Equation 8 follows [16]:

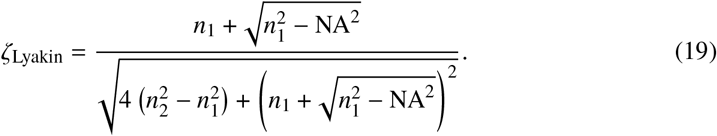

The same value was proposed earlier by Stallinga while posing different criteria of minimal variation of the phase function: *θ*^*∗*^ : min ⟨ (Φ − Φ(*θ*^*∗*^))^2^ ⟩ [15]. For Diel *et al*. the ζ_median_ corresponds to setting 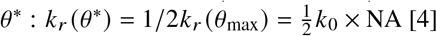, such that

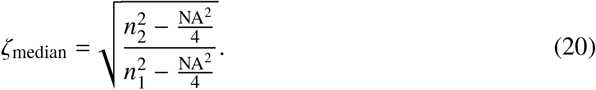

These estimates for ζ are geometrical in nature, where the actual wavelength is not taken into account – with the radii of the Ewalds’ spheres being cancelled out of the fraction in Equation 8. This results in an implicit omission of the interference nature of the PSF under RIM. Somewhat less obviously, the criterion of Stallinga is also geometrical [15]. While it does demand minimal variation in the value of Φ in order to achieve the highest possible constructive interference, it does not take into account that the well minimized phase function can become several 2π in range (when both *z* _*A*_ and *z*_*N*_ are large enough) and thus lead to destructive interference anyway. In order to be able to correct for depth-dependent axial scaling in experiments where imaging under RIM is unavoidable, it is crucial that (i) the position of the coverslip (or more general, the RIM interface) is determined precisely and, (ii) spherical aberration at the interface is reduced to an absolute minimum by selecting the cover glass the objective is optimized for, or by optimizing the position of the correction collar. In practical terms, this would require that the objective lens used is fitted with a correction collar, which is often the case for high-NA objective lenses.

One such case would be cryo-fluorescence microscopy used in the cryo-electron tomography workflow, where a RIM is generally present due to the large temperature difference between the optical objective and the sample. With fluorescence microscopy, targets can be identified in frozen hydrated cells and consequently sufficiently thin sections can be prepared by ablating the excess cellular material surrounding the target with a focused ion beam [37–39]. The aimed thickness of this frozen section is approximately 100 to 200 nm, which makes it crucial to precisely determine the target position with respect to the RIM interface (outer cell surface) [40]. If axially scaled distances measured with the light microscope are not corrected, targeting errors in the range of 300 to 1200 nm will occur with cell thicknesses ranging up to micrometers. Moreover, the *depth-dependence* of these errors will be important when fabricating sections out of thicker ice whilst using fluorescence microscopy to find targets in for instance organoids, as can be done in high pressure frozen samples [41, 42].

With the provided software [32], one can easily plot the depth-dependent re-scaling factor for the relevant imaging scenario, but also re-scale their 3D microscopy data, without having to judge which linear theory will hold in this scenario or perform full wave-optics calculations. We should note, however, that although the data will be axially re-scaled, our software does not provide any correction of the shape of the point spread function (PSF) due to spherical aberrations. To correct for this, one would require deconvolution with a depth-dependent PSF as demonstrated in [43]. Our measurements are limited by the relatively high uncertainty in determining the re-scaling factor near *z* _*A*_ = 0. It would therefore be interesting, for future research, to use the method from Petrov *et al*. to measure the re-scaling factor for the scenario with a 0.85 NA air objective when imaging *n*_1_ = 1.0 → *n*_2_ = 1.33.

## 6. Conclusion

We have presented an analytical theory to correct for the depth-dependent axial deformation when imaging with light microscopy in the presence of a refractive index mismatch between the sample and the microscope objective immersion medium. Using the resulting equation, a re-scaling factor as a function of depth can be calculated from imaging parameters numerical aperture (NA), the refractive indices of the objective (*n*_1_) and sample (*n*_2_), and the wavelength (*λ*), for a RIM with both *n*_1_ < *n*_2_ and *n*_1_ > *n*_2_. We performed wave-optics calculations to verify the theory and find a very good agreement between the two. In addition, we performed experiments to measure the axial scaling, where we find a good agreement at larger depths, whereas closer to the cover slip, the measurements suffer from large uncertainties. We do find good agreement with the accurate measurements done by Petrov *et al*. [23] in the case when imaging with a high-NA oil immersion objective into a water sample.

Next, we tested the depth-dependent theory versus existing linear theories in the literature on 3D microscopy data with a known ground truth. We find that for NA = 0.85, *n*_1_ = 1, *n*_2_ = 1.33 the depth-dependent theory outperforms existing linear theories for depths up to 80 µm with a maximum relative error of 2 %. For NA = 1.4, *n*_1_ = 1.52, *n*_2_ = 1.33, the depth-dependent theory performs as good as the best linear theory in literature. However, we think its performance was compromised by the presence of a small spherical aberration near the cover slip, which could not be corrected for as the objective did not have a correction collar. Moreover, we could not compare close to the cover glass (< 10 µm), where the linear theory used in the comparison was expected to break down.

Finally, we have presented a web applet to be used by microscope users to calculate the re-scaling factor for their imaging parameters. In addition, we have shared software to re-scale 3D data sets using the depth-dependent scaling factor.

Our re-scaling theory is the first to include the depth dependence of axial scaling due to a refractive index mismatch. It will be of use in imaging scenarios where the refractive indices of the sample and objective cannot be matched, such as in integrated, cryogenic, and correlative light and electron microscopy setups, or in the imaging of water-like samples using high-NA oil immersion objectives.

## Funding

This work is part of the Cryo3Beams project (project N° 17152) financed via the High Tech Systems and Materials programme of the Applied and Engineering Sciences domain of the Dutch Research Council (NWO).

## Acknowledgments

We thank Hans C. Gerritsen and Gerhard A. Blab for the useful discussions, Bernd Rieger for feedback on our results and Alfons van Blaaderen for the use of the refractometer.

## Disclosures

SVL, MNFH and EBW declare no conflicts of interest. DBB is employed by Delmic BV, JPH has a financial interest in Delmic B.V.

## Data availability

Data underlying the results presented in this paper are available in Ref. [44].

## Supplemental document

See Supplement 1 for supporting content.

## Notes

### Summary of Updates

Figures have been merged and/or moved to the supplemental information for readability purposes. Some mistakes in the derivation have been corrected. Additional text is added to clarify ambiguities

