## Supplement 1 for "Depth-dependent scaling of axial distances in light microscopy"

### Depth-dependent scaling of axial distances in light microscopy: supplemental document

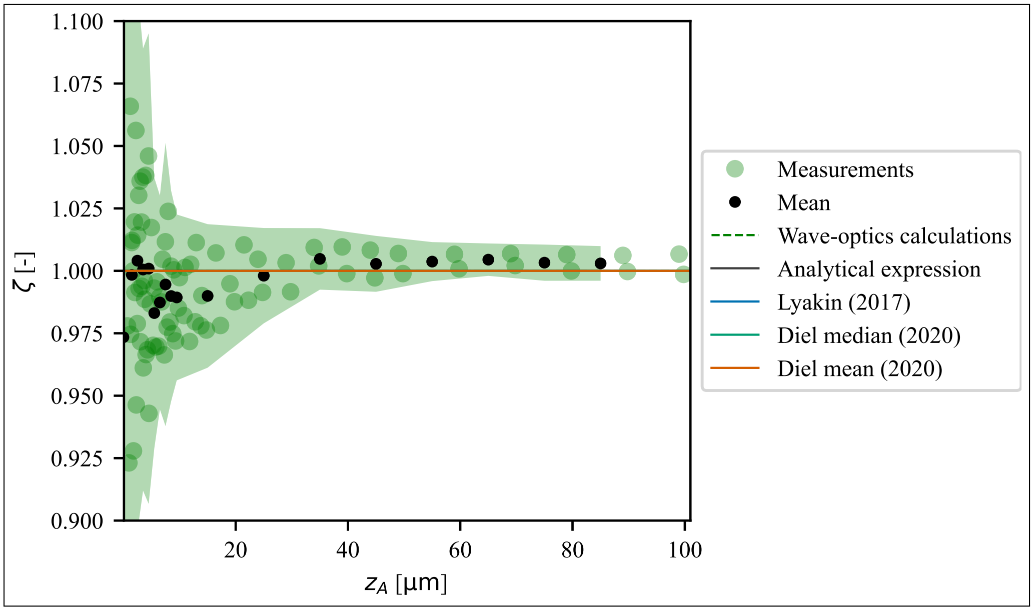


**Fig. S1.** Re-scaling factor versus depth for 0.7 NA optical objective, $n_1 = 1.0$ and $n_2 = 1.0$. Measurement data (green dots) are plotted alongside the wave-optics calculations (dashed blue lines), analytical solution (solid black line) and depth-independent theories (solid lines). The wave-optics calculations, analytical solution and depth-independent theories overlap at $\zeta = 1.0$. 2 sets of measurement data are plotted individually and from this data the mean (solid black dots) is computed by binning along $z_A$, plotted in the center of each bin (bin sizes of 1 and 10 µm respectively for 0 to 10 µm and 10 to 100 µm). The measurement error (shaded green area) is the sum of the measurements standard deviation and an estimated upper error limit.

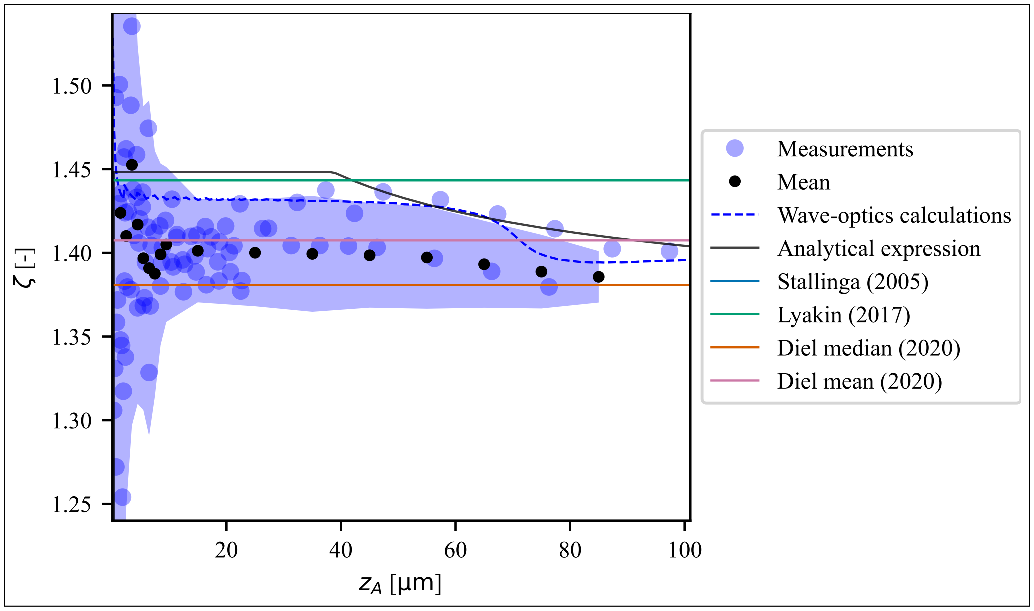


**Fig. S2.** Re-scaling factor versus depth for 0.7 NA optical objective, $n_1 = 1.0$ and $n_2 = 1.34$. Measurement data (blue dots) are plotted alongside the wave-optics calculations (dashed blue lines), analytical solution (solid black line) and depth-independent theories (solid lines). 4 sets of measurement data are plotted individually and from this data the mean (solid black dots) is computed by binning along $z_A$, plotted in the center of each bin (bin sizes of 1 and 10 µm respectively for 0 to 10 µm and 10 to 100 µm). The measurement error (shaded blue area) is the sum of the measurements standard deviation and an estimated upper error limit.

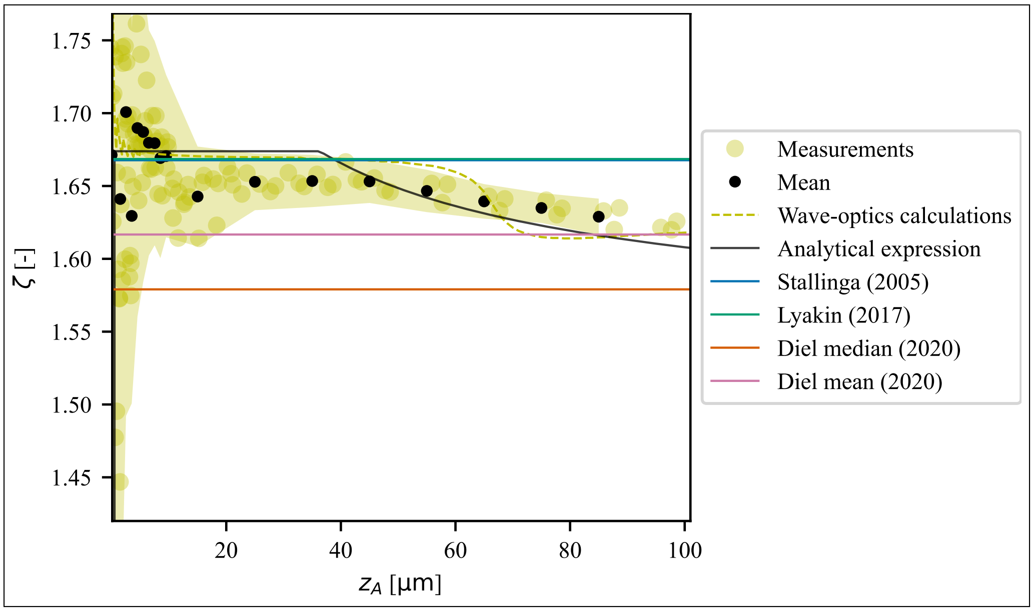


**Fig. S3.** Re-scaling factor versus depth for 0.7 NA optical objective, $n_1 = 1.0$ and $n_2 = 1.52$. Measurement data (yellow dots) are plotted alongside the wave-optics calculations (dashed blue lines), analytical solution (solid black line) and depth-independent theories (solid lines). 3 sets of measurement data are plotted individually and from this data the mean (solid black dots) is computed by binning along $z_A$, plotted in the center of each bin (bin sizes of 1 and 10 µm respectively for 0 to 10 µm and 10 to 100 µm). The measurement error (shaded yellow area) is the sum of the measurements standard deviation and an estimated upper error limit.

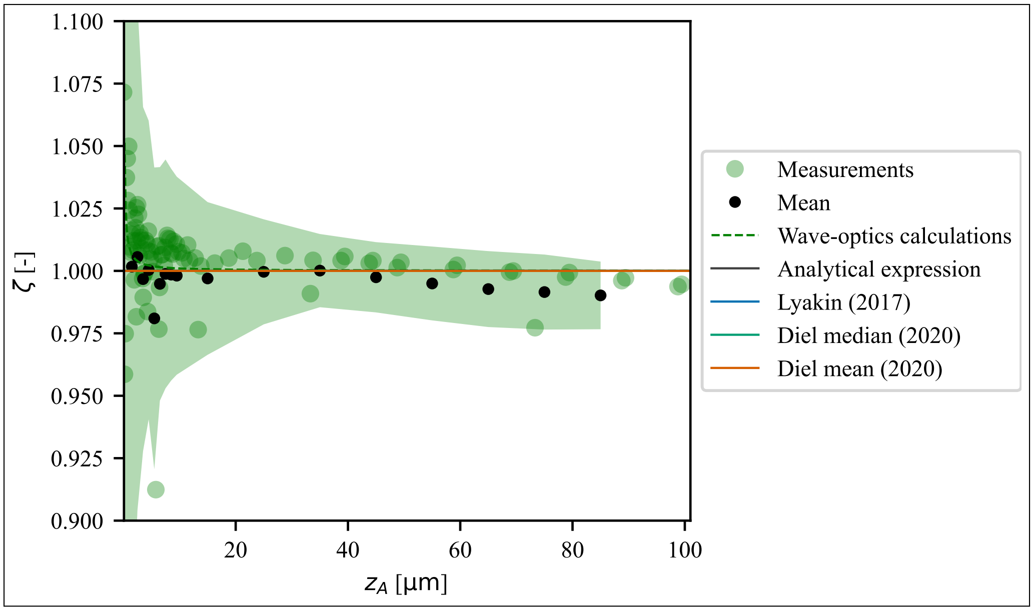


**Fig. S4.** Re-scaling factor versus depth for 0.85 NA optical objective, $n_1 = 1.0$ and $n_2 = 1.0$. Measurement data (green dots) are plotted alongside the wave-optics calculations (dashed blue lines), analytical solution (solid black line) and depth-independent theories (solid lines). The wave-optics calculations, analytical solution and depth-independent theories overlap at $\zeta = 1.0$. 3 sets of measurement data are plotted individually and from this data the mean (solid black dots) is computed by binning along $z_A$, plotted in the center of each bin (bin sizes of 1 and 10 µm respectively for 0 to 10 µm and 10 to 100 µm). The measurement error (shaded green area) is the sum of the measurements standard deviation and an estimated upper error limit.

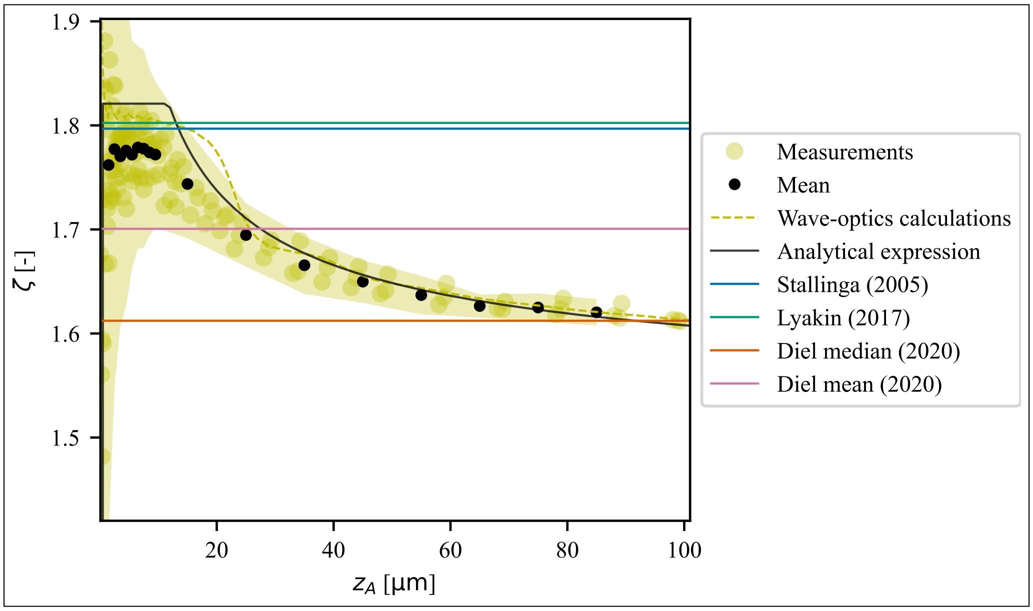


**Fig. S5.** Re-scaling factor versus depth for 0.85 NA optical objective, $n_1 = 1.0$ and $n_2 = 1.52$. Measurement data (yellow dots) are plotted alongside the wave-optics calculations (dashed blue lines), analytical solution (solid black line) and depth-independent theories (solid lines). 3 sets of measurement data are plotted individually and from this data the mean (solid black dots) is computed by binning along $z_A$, plotted in the center of each bin (bin sizes of 1 and 10 µm respectively for 0 to 10 µm and 10 to 100 µm). The measurement error (shaded yellow area) is the sum of the measurements standard deviation and an estimated upper error limit.

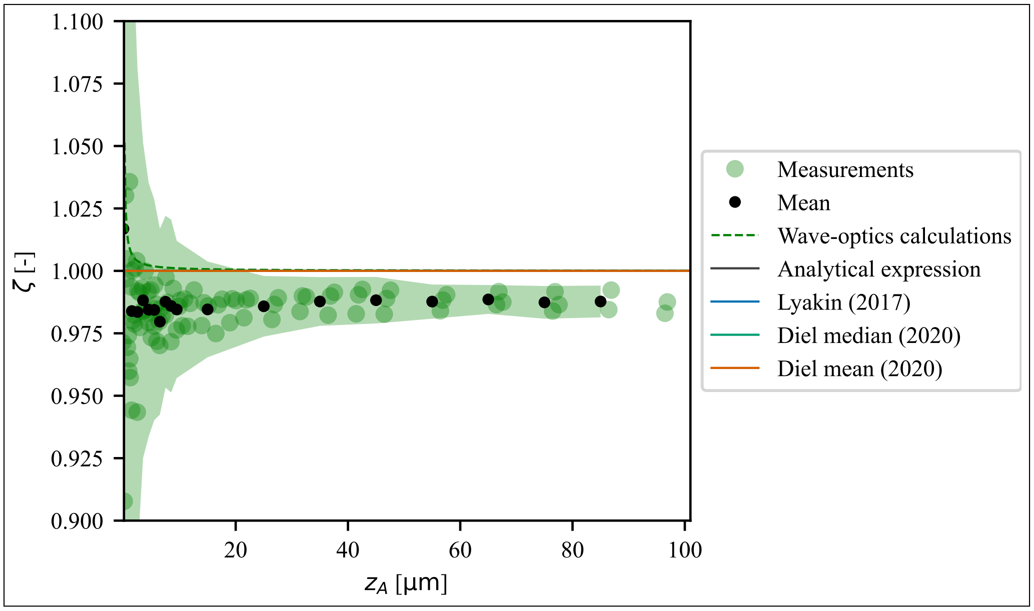


**Fig. S6.** Re-scaling factor versus depth for 0.95 NA optical objective, $n_1 = 1.0$ and $n_2 = 1.0$. Measurement data (green dots) are plotted alongside the wave-optics calculations (dashed blue lines), analytical solution (solid black line) and depth-independent theories (solid lines). The wave-optics calculations, analytical solution and depth-independent theories overlap at $\zeta = 1.0$. 3 sets of measurement data are plotted individually and from this data the mean (solid black dots) is computed by binning along $z_A$, plotted in the center of each bin (bin sizes of 1 and 10 µm respectively for 0 to 10 µm and 10 to 100 µm). The measurement error (shaded green area) is the sum of the measurements standard deviation and an estimated upper error limit.

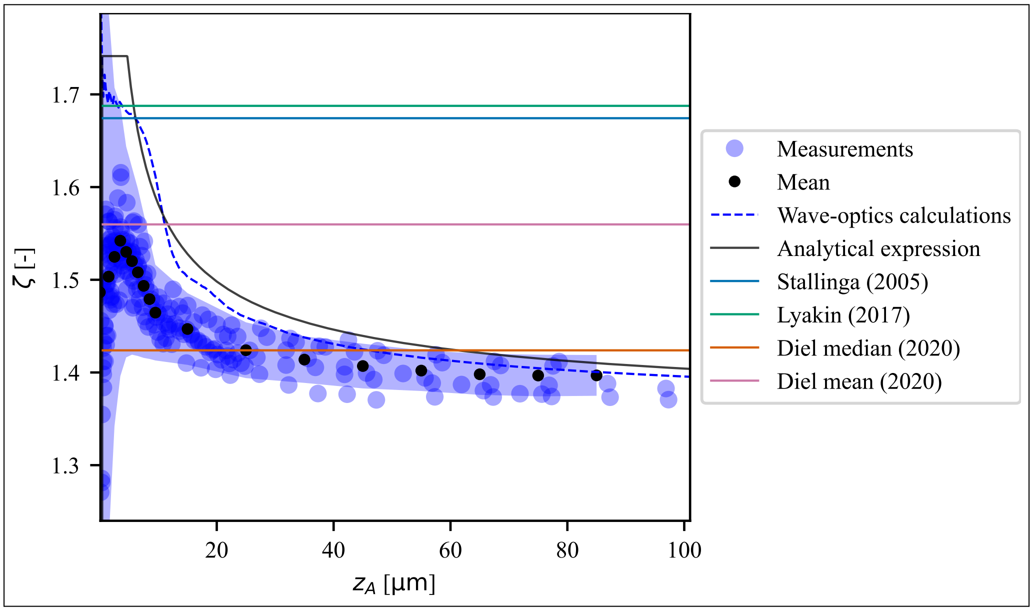


**Fig. S7.** Re-scaling factor versus depth for 0.95 NA optical objective, $n_1 = 1.0$ and $n_2 = 1.34$. Measurement data (blue dots) are plotted alongside the wave-optics calculations (dashed blue lines), analytical solution (solid black line) and depth-independent theories (solid lines). 9 sets of measurement data are plotted individually and from this data the mean (solid black dots) is computed by binning along $z_A$, plotted in the center of each bin (bin sizes of 1 and 10 µm respectively for 0 to 10 µm and 10 to 100 µm). The measurement error (shaded blue area) is the sum of the measurements standard deviation and an estimated upper error limit.

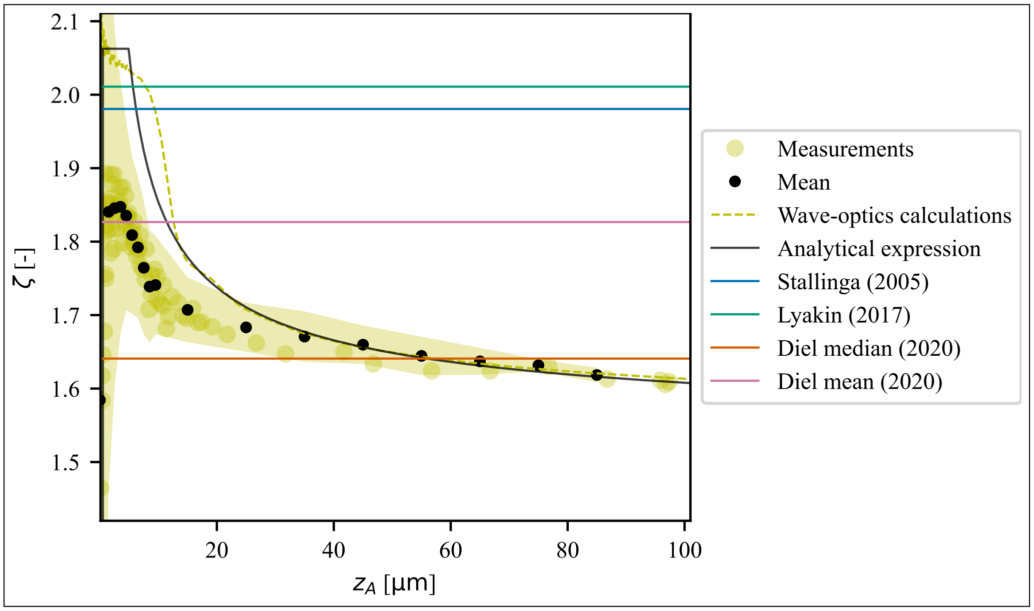


**Fig. S8.** Re-scaling factor versus depth for 0.95 NA optical objective, $n_1 = 1.0$ and $n_2 = 1.52$. Measurement data (yellow dots) are plotted alongside the wave-optics calculations (dashed blue lines), analytical solution (solid black line) and depth-independent theories (solid lines). 3 sets of measurement data are plotted individually and from this data the mean (solid black dots) is computed by binning along $z_A$, plotted in the center of each bin (bin sizes of 1 and 10 µm respectively for 0 to 10 µm and 10 to 100 µm). The measurement error (shaded yellow area) is the sum of the measurements standard deviation and an estimated upper error limit.

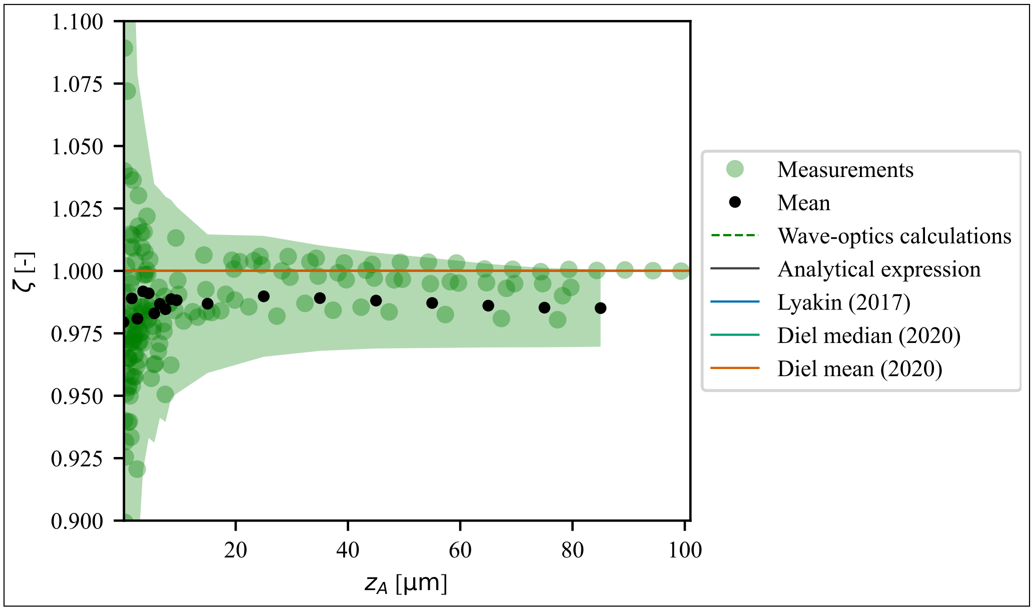


**Fig. S9.** Re-scaling factor versus depth for 1.25 NA optical objective, $n_1 = 1.34$ and $n_2 = 1.34$. Measurement data (green dots) are plotted alongside the wave-optics calculations (dashed blue lines), analytical solution (solid black line) and depth-independent theories (solid lines). The wave-optics calculations, analytical solution and depth-independent theories overlap at $\zeta = 1.0$. 6 sets of measurement data are plotted individually and from this data the mean (solid black dots) is computed by binning along $z_A$, plotted in the center of each bin (bin sizes of 1 and 10 µm respectively for 0 to 10 µm and 10 to 100 µm). The measurement error (shaded green area) is the sum of the measurements standard deviation and an estimated upper error limit.

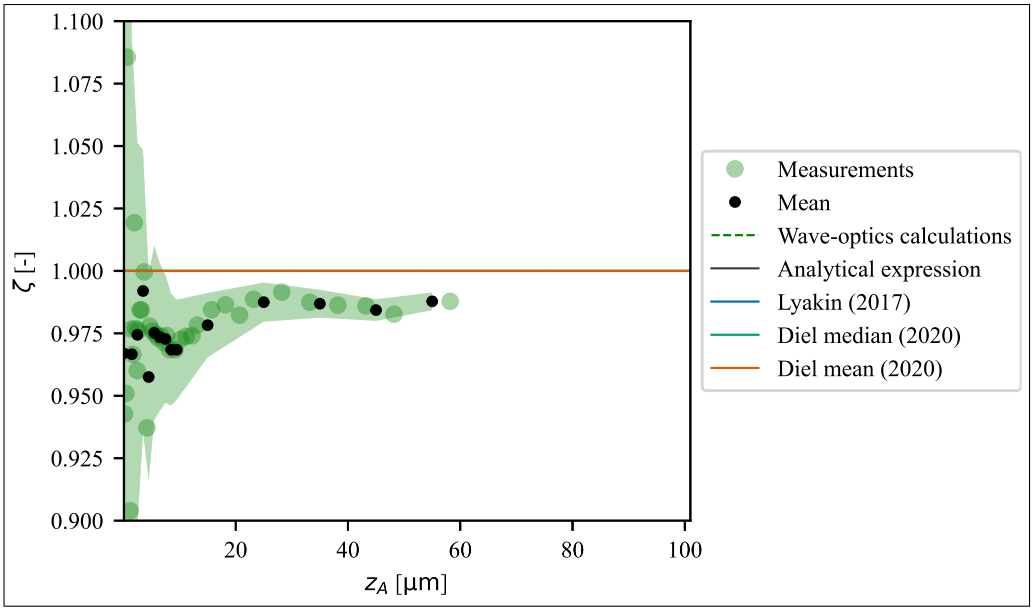


**Fig. S10.** Re-scaling factor versus depth for 1.4 NA optical objective, $n_1 = 1.52$ and $n_2 = 1.52$. Measurement data (green dots) are plotted alongside the wave-optics calculations (dashed blue lines), analytical solution (solid black line) and depth-independent theories (solid lines). The wave-optics calculations, analytical solution and depth-independent theories overlap at $\zeta = 1.0$. 1 sets of measurement data are plotted individually and from this data the mean (solid black dots) is computed by binning along $z_A$, plotted in the center of each bin (bin sizes of 1 and 10 μm respectively for 0 to 10 μm and 10 to 100 μm). The measurement error (shaded green area) is the sum of the measurements standard deviation and an estimated upper error limit.

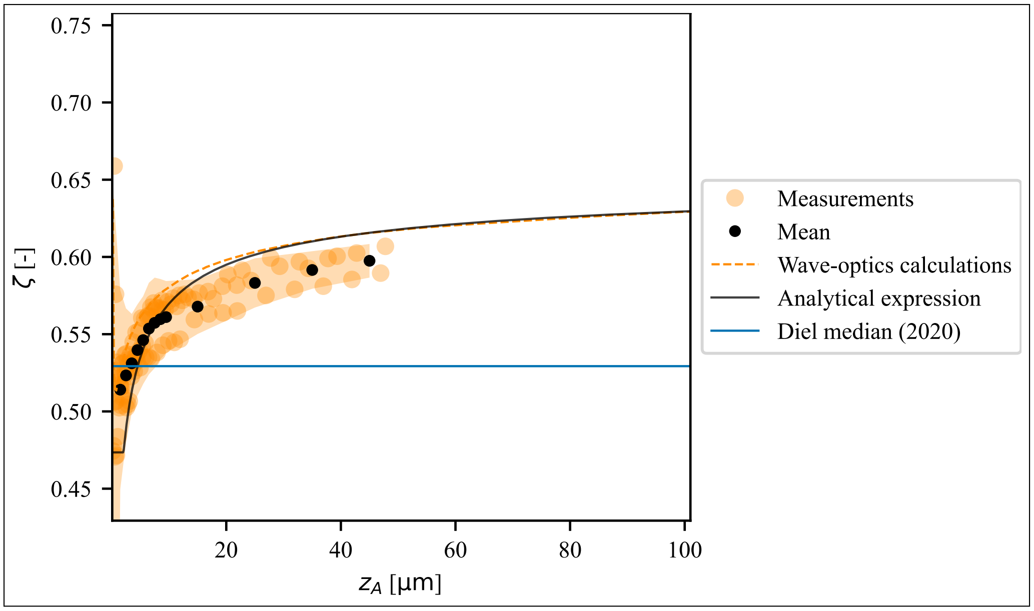


**Fig. S11.** Re-scaling factor versus depth for 1.4 NA optical objective, $n_1 = 1.52$ and $n_2 = 1.0$. Measurement data (dark orange dots) are plotted alongside the wave-optics calculations (dashed blue lines), analytical solution (solid black line) and depth-independent theories (solid lines). 3 sets of measurement data are plotted individually and from this data the mean (solid black dots) is computed by binning along $z_A$, plotted in the center of each bin (bin sizes of 1 and 10 µm respectively for 0 to 10 µm and 10 to 100 µm). The measurement error (shaded dark orange area) is the sum of the measurements standard deviation and an estimated upper error limit.

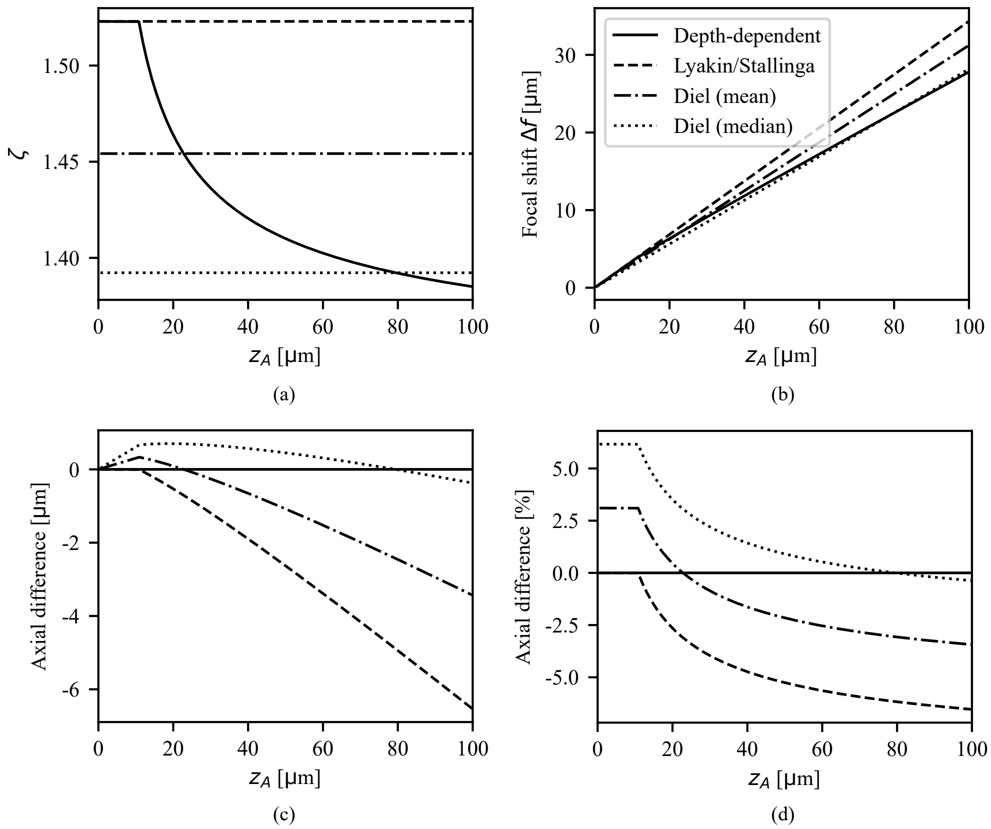


**Fig. S12.** Quantitative comparison of the linear scaling theories from literature to the depth-dependent analytical theory for NA=0.85, $n_1 = 1.0$ and $n_2 = 1.336$. **(a)** The re-scaling factor $\zeta$ vs AFP for the depth-dependent theory (solid), Lyakin *et al.* (dashed) [1], Diel *et al.* mean (dash-dot) and median (dot-dot) [2]. **(b)** The focal shift vs AFP of the aforementioned theories. **(d)** Axial difference vs AFP for the theories. **(d)** Relative difference between the theories.

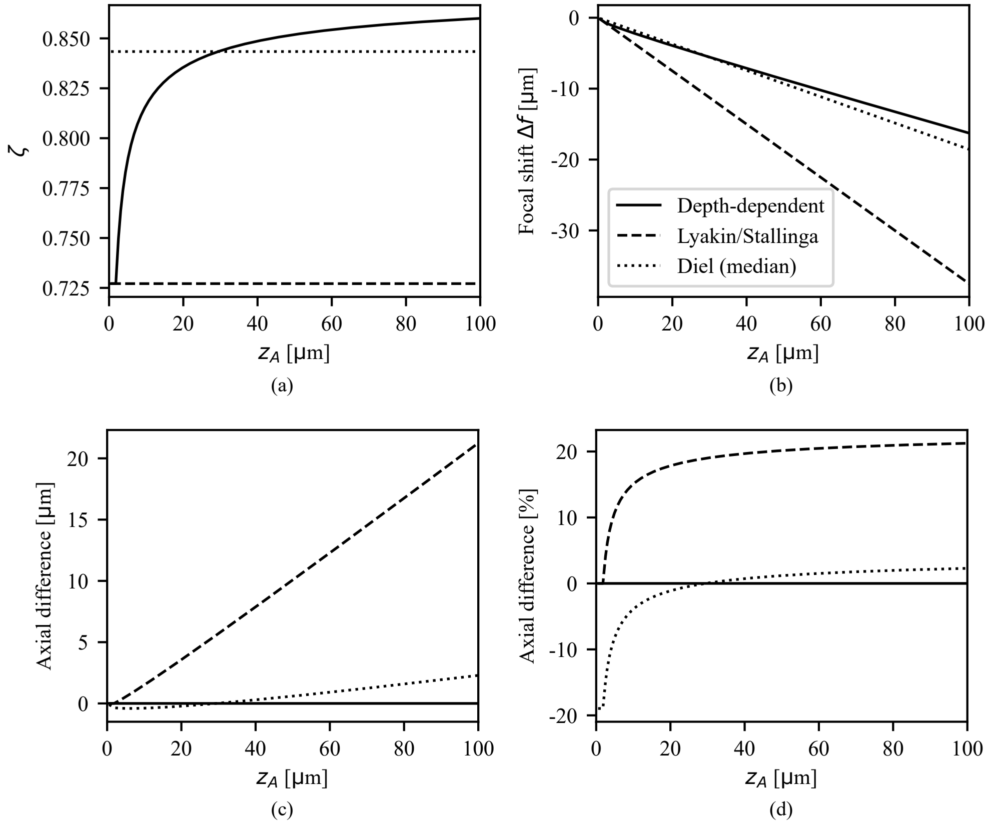


**Fig. S13.** Quantitative comparison of scaling theories in case of NA = 1.4, $n_1 = 1.52$, $n_2 = 1.336$. **(a)** The re-scaling factor $\zeta$ vs AFP for the depth-dependent theory (solid), Lyakin *et al.* (dashed) [1] and median (dot-dot) [2]. **(b)** The focal shift vs AFP of the aforementioned theories. **(d)** Axial difference vs AFP for the theories. **(d)** Relative difference between the theories.
